# Association of a Common Genetic Variant with Parkinson’s Disease is Propagated through Microglia

**DOI:** 10.1101/2021.01.15.426824

**Authors:** R. G. Langston, A. Beilina, X. Reed, A. B. Singleton, C. Blauwendraat, J. R. Gibbs, M. R. Cookson

**Author notes:** Correspondence to: Mark R. Cookson PhD.

## Abstract

Studies of the genetic basis of Parkinson’s disease (PD) have identified many disease-associated genetic variants, but the mechanisms linking variants to pathogenicity are largely unknown. PD risk is attributed to both coding mutations in the Leucine-rich repeat kinase 2 (*LRRK2*) gene and to common non-coding variation upstream of the *LRRK2* locus. Here we show that the influence of genotype at non-coding variant rs76904798 on *LRRK2* expression is propagated specifically through microglia, in contrast to evaluations based on general rather than genotype-dependent expression. We find evidence of microglia-specific regulatory regions that may modulate *LRRK2* expression using single nuclei sequencing analyses of human frontal cortex and confirm these results in a human induced pluripotent stem cell-derived microglia model. Our study demonstrates that cell type is an important consideration in interrogation of the role of non-coding variation in disease pathogenesis.

## Introduction

Genome-wide association studies (GWAS) have been increasingly used to identify genetic contributions to human diseases. In most disease GWAS designs, the differences in frequency of common single nucleotide polymorphisms (SNPs) between cases and controls are used to infer which genomic regions influence lifetime disease risk. However, as linkage disequilibrium (LD) within the human genome results in the co-inheritance of a series of SNPs, it is difficult to infer from GWAS which gene or variants are causally associated with disease risk. An exception to this general rubric is where the GWAS locus contains a gene independently shown to contribute to disease risk by virtue of familial inheritance for the same condition.

Such polygenic risk loci include a locus on human chromosome 12 that includes the gene encoding Leucine-rich repeat kinase 2 (*LRRK2*). Coding variants in *LRRK2* are causal for familial Parkinson’s disease (PD) while non-coding variants in the 5’ region of the gene have been nominated by GWAS as increasing sporadic PD risk (*1–3*). Thus, we can be confident that *LRRK2* is the causal gene at this locus.

Because of the inherent limitations of GWAS, there have been many approaches that attempt to prioritize candidate genes and variants using additional datasets. Approaches that nominate enrichment in specific cell types have been useful in identifying likely genes as most tissues have heterogeneous cellular composition. Several approaches have nominated dopamine neurons (*4–7*), which are principal cell types lost in PD, as mediators of neurodegeneration, although other studies have identified non-neuronal cells including oligodendrocytes and oligodendrocyte precursor cells (*4, 8*) as contributing to disease risk. Such approaches rely on the presumed mechanism that non-coding variants affect gene expression, which can be inferred by mapping expression quantitative trait loci (eQTL). It has been shown previously that there is a *LRRK2* eQTL in peripheral monocytes and in microglia-like cells derived from peripheral B cells (*9*). However, *LRRK2* is also strongly expressed in excitatory neurons in the cerebral cortex (*10–12*) and spiny projection neurons in the striatum (*13–16*), the latter being targets of nigral dopamine neurons. Finally, *LRRK2* is also expressed in oligodendrocyte precursor cells (*10*). It is therefore unclear which cells in the human brain mediate PD risk generally and *LRRK2* expression as a quantitative trait specifically.

Here, we experimentally determine which cells in the human brain contribute to the disease-linked expression as a quantitative trait at the *LRRK2* locus. A summary of our experimental workflow is presented in Fig. 1A. We found evidence of genotype-dependent *LRRK2* expression only in microglia despite robust expression across multiple cell types, including excitatory neurons and oligodendrocyte precursor cells as previously documented. Furthermore, we could recapitulate the same genotype to gene expression relationship in iPSC-derived microglia from donors with risk or protective alleles. Finally, we map microglial-specific areas of open chromatin that we nominate as a likely mechanistic explanation for the observed differences in genotype-dependent *LRRK2* expression between cell types. These results show that interpretation of GWAS results can be improved by examining the relationship between genotype and cell type-dependent gene expression.

**Fig. 1.**
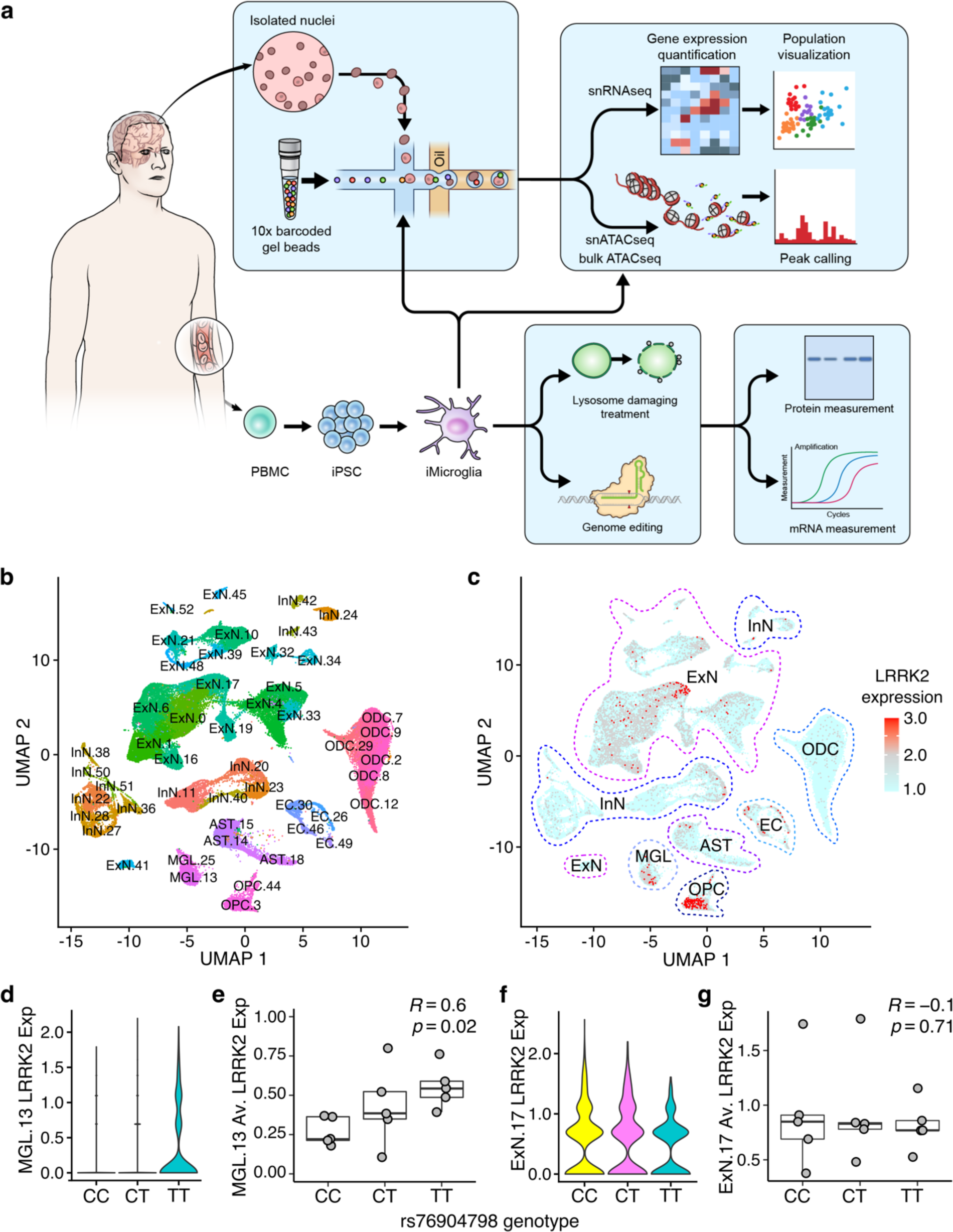
Genotype at rs76904798 has a microglia-specific influence on *LRRK2* expression in human frontal cortex. (A) Schematic depiction of experimental workflow. Nuclei isolated from human frontal cortex were subjected to single nuclei RNAseq (snRNAseq) and single nuclei ATACseq (snATACseq) analyses. Induced pluripotent stem cells (iPSC) generated from patient-derived peripheral blood mononuclear cells (PBMC) were differentiated to microglia-like cells (iMicroglia) and evaluated by single cell RNA sequencing, bulk ATAC sequencing, and western blot and qPCR assays following perturbations including a lysosome damaging treatment and CRISPR/Cas-directed genome editing. Graphic design credit to Erina He at NIH Medical Arts. (B) UMAP visualization of cell clusters identified in an integrated snRNAseq dataset derived from 15 human frontal cortex samples, labeled based on differential expression of known cell type marker genes. A total of 113,520 nuclei are represented. (C) Localization of *LRRK2* expression in brain cell populations, grouped according to broad cell type (excitatory neuron, ExN; inhibitory neuron, InN; oligodendrocyte, ODC; oligodendrocyte precursor cell, OPC; endothelial cell, EC; astrocyte, AST; microglia, MGL) and identified by relative abundance of that cell population in the overall dataset. (D) Violin plot showing the distribution of *LRRK2* expression in cell population MGL.13 split by genotype at rs76904798. (E) Boxplot where each point represents average *LRRK2* expression of cells in MGL.13 contributed by each of the 15 donors. Using the number of minor alleles (CC = 0, CT = 1, TT = 2) as the x-axis in a simple linear regression model, Pearson’s *R* = 0.598 (*p* = 0.019, *t* = 2.69, df = 13). Plots in (F) and (G) show the equivalent analyses for cell population ExN.17, Pearson’s *R* = −0.103 (*p* = 0.714, *t* = −0.37, df = 13).

## Results

### The rs76904798:LRRK2 eQTL is expressed specifically in microglia in human brain

As *LRRK2* is expressed in some neuronal subtypes and in non-neuronal cells, we first evaluated which cell type mediates the *LRRK2* eQTL associated with PD in an unbiased manner.

To achieve this, we performed single nuclei RNA sequencing on brain samples from fifteen donors representing the three genotypes (CC, CT and TT) at the lead SNP for PD risk, rs76904798[C/T]. A total of 117,632 nuclei were captured with an average sequencing depth of 27,000 reads per nucleus. There were no significant differences between genotype groups in the number of nuclei measured, the mean reads per nucleus, the median number of genes per cell or total number of genes detected (Fig. S1A-D). Following integration and normalization of the 15 datasets, 54 cell populations were identified using shared-nearest-neighbor clustering. Clusters were assigned a cell type based on differential expression of known cell type marker genes (*11, 17–19*) (Fig. S1E). Five clusters were found to express markers of multiple cell types and were excluded from downstream analysis. The remaining clusters (113,520 nuclei) were labeled with their broad cell type (excitatory neuron, ExN; inhibitory neuron, InN; endothelial cell, EC; oligodendrocyte precursor cell, OPC; oligodendrocyte, ODC; astrocyte, AST; microglia, MGL) and their original cluster number to distinguish cell subtype populations, with cluster number indicating rank of each cluster according to size (Fig. 1B). These clusters contained nuclei from every sample, except for cluster ODC.29 (an oligodendrocyte cluster that is the 29th most populous of all cell populations) which contained nuclei from all donors except donor S1135 (Fig. S1F). The proportion of each cluster contributed by donors of each rs76904798 genotype group is shown in Fig. S1G. These results demonstrate that we were able to successfully identify all expected brain cell types via snRNAseq of post-mortem human frontal cortex, with equivalent recovery of these cell types between donors.

We next examined which cell types expressed *LRRK2* in the same samples (Fig. 1C, Fig. S2A). *LRRK2* expression was notably high in oligodendrocyte precursor cell population OPC.3, excitatory neuron population ExN.17 and microglia population MGL.13, i.e., a cluster of cells positively identified as microglia that is the 13th most numerous of all cells in this experiment. We validated the same pattern of cell-type dependent *LRRK2* expression in the human substantia nigra using publically available data, showing that expression could be readily detected in oligodendrocyte precursor cells and microglia (*4, 20*) (Fig. S3). These results show that multiple cell types in both human frontal cortex and substantia nigra express *LRRK2*. We then tested for any cell type-specific effect of rs76904798 genotype on *LRRK2* expression by performing differential expression analyses of genotype groups in each cell population. A significant difference in *LRRK2* expression between genotypes was observed in the microglia population MGL.13, with differentially higher expression of *LRRK2* in cells homozygous for the risk variant (TT cells) compared to CC cells (adjusted *p* value = 7.24×10^-18^, non-parametric Wilcoxon rank sum test followed by Bonferroni multiple test correction; *N* = 5 frontal cortex samples per genotype; *n*CC = 1460, *n*TT = 704 nuclei; average log2(fold change) = −0.26 per T allele; Fig. 1D). Of all other cell populations, *LRRK2* expression was also significantly different between CC and TT cells of cluster EC.46, but with the opposite direction of effect with higher expression in CC cells compared to TT cells (Fig. S2B; adjusted *p* value = 3.61 ×10^-5^; *N* = 5 frontal cortex samples per genotype; *n*CC = 330, *n*TT = 183 nuclei; average log2(fold change) = 0.33 per T allele). We also assessed if dosage of the T allele was correlated with *LRRK2* expression in any cell type. Average *LRRK2* expression in each cluster was calculated by donor and linear regression indicated a significant correlation between expression of *LRRK2* and the presence of the T variant at rs76904798 only in microglia population MGL.13 (Fig. 1E; Pearson’s *R* = 0.60, *p* = 0.019, *n* = 15 frontal cortex samples). No relationship between rs76904798 genotype and *LRRK2* expression was identified in excitatory neuron population ExN.17, in which *LRRK2* is highly expressed, which excludes that lack of a relationship in this cell type was driven by low expression levels (Fig. 1F-G). Extending these tests to all cell populations, no other genotype-dependent differences in *LRRK2* expression were detected other than in population MGL.13 (plots shown for populations in which *LRRK2* was detected are shown in Fig. S2B). Therefore, in the human brain we find a microglial correlation between rs76904798 genotype and *LRRK2* expression, identifying a eQTL for PD risk that is stronger in microglia than in other cell types. *iPSC-derived microglia are transcriptionally similar to brain microglia and respond to stimulation* We next looked for an experimentally tractable model that would allow us to confirm or refute the eQTL analysis and to also examine the relationship between genotype and LRRK2 kinase activity. We confirmed that LRRK2 was robustly expressed in iPSC-derived microglia (iMicroglia) (*21, 22*) by western blot analysis of iMicroglia differentiated from control iPSC alongside LRRK2 knockout cells generated by CRISPR/Cas9-directed genome editing (Fig. S4).

We then differentiated iPSC lines from the Parkinson’s Progression Marker Initiative (PPMI) series to iMicroglia. As it was important that differentiated cells were transcriptionally similar to human brain microglia, we first characterized their gene expression characteristics. iMicroglia derived from two donors were submitted to single cell RNA sequencing (scRNAseq). Clustering analysis of these data revealed six sub-populations (Fig. S5A), all expressing *LRRK2* (Fig. S5B) and microglia marker genes (Fig. S5C). Importantly, the cell lines differentiated consistently, with iMicroglia derived from both lines contributing to each cluster. Average gene expression between cells of each line in each cluster was highly correlated, with Pearson’s *R* = 0.97 (*p* < 2.3 ×10^-16^, *n* = 17,108 features) in every cluster except cluster “iMGL_5” (Pearson’s *R* = 0.88, *p* < 2.3 ×10^-16^, *n* = 17,108 features; Fig. S5D). To directly compare the iPSC-derived microglia to human brain cells, the iMicroglia scRNAseq dataset was integrated with the snRNAseq dataset generated from the human frontal cortex. The iMicroglia mapped to brain glial cells on a UMAP dimensionality reduction plot (Fig. 2A) and their overall pattern of gene expression was most highly correlated with brain microglia (Pearson’s *R* = 0.66, *p* < 2.3 ×10^-16^, *n* = 31,133 features) compared to other brain cell types (Fig. 2B). These results show that iPSC-derived microglia are transcriptionally similar to microglia of the human brain.

**Fig. 2.**
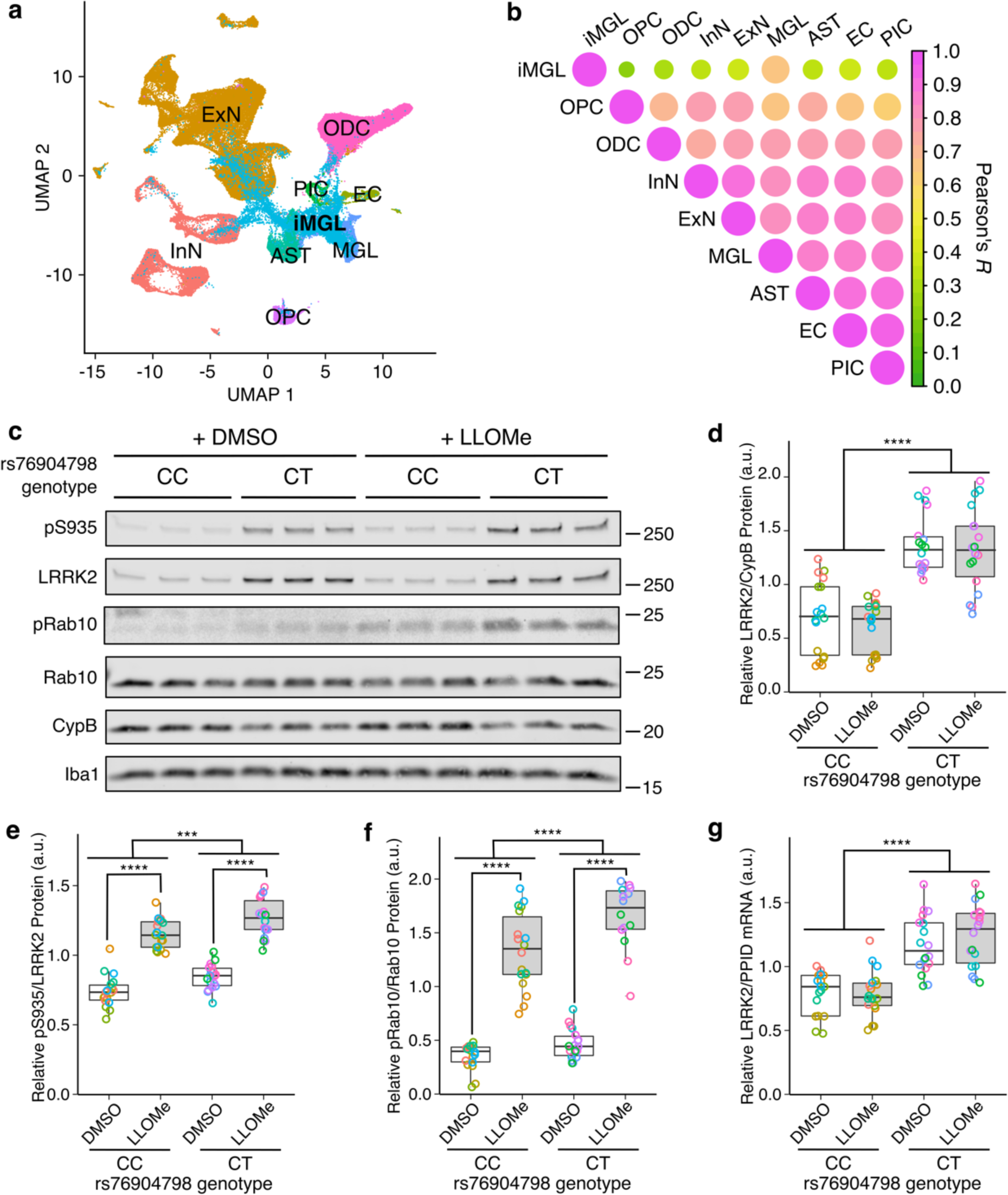
iMicroglia carrying PD risk variant rs76904798-T have increased LRRK2 protein and activity levels. (A) UMAP visualization of the iMicroglia scRNAseq dataset integrated with the human frontal cortex snRNAseq dataset, with clusters labeled according to broad cell type (excitatory neuron, ExN; inhibitory neuron, InN; oligodendrocyte, ODC; oligodendrocyte precursor cell, OPC; endothelial cell, EC; astrocyte, AST; microglia, MGL; peripheral immune cell, PIC; iPSC-derived microglia, iMGL. (B) Spot matrix illustrating the correlation between average expression of 31,133 transcripts between the iMGL population and brain cell type populations, described by Pearson’s correlation coefficient. For iMGL vs MGL, *R* = 0.658, df = 31,131, *p* = 2.22 x 10^-16^, *t* = 154.17). (C) Representative western blot analysis of a pair of differentiated iMicroglia, PPMI 3411 (rs76904798-CC) and PPMI 3953 (rs76904798-CT), with immunoblotting for phospho-Ser^935^-LRRK2, total LRRK2, phospho-Rab10, total Rab10, cyclophilin b (CypB) and Iba1 -/+ 15-minute LLOMe treatment. (D-G) Quantifications of normalized mRNA or protein levels, shown relative to the mean values for each of six experiments. The *p*(genotype) results of two-way ANOVA are shown for each measurement, *N* = 6 cell lines per genotype, *n* = 17-18 per genotype-treatment group with technical replicates. The post-hoc analysis results of Tukey’s multiple comparisons test between treatment groups are shown where statistically significant. *P<0.05, **P<0.01, ***P<0.001, ****P<0.0001.

In this cell model, we could measure LRRK2 kinase activity based on the phosphorylation of physiological substrate Rab10 (*23*). As well as baseline activity, we explored methods of microglia stimulation and LRRK2 activation by exposing cells for two hours to lipopolysaccharide (LPS), α-synuclein conditioned medium (aSCM), or L-leucyl-L-leucine methyl ester (LLOMe), which we have recently shown to activate LRRK2 in primary mouse astrocytes (*24*). Immunoblot analysis indicated that of the three treatments, LLOMe most strongly activated LRRK2 kinase activity in iMicroglia as measured by significantly increased phosphorylation of Rab10 (Fig. S6A-E). Induction of pro-inflammatory stress-response pathways was observed following exposure to all three treatments, with LPS and aSCM initiating upregulation of NF-κB signaling while LLOME induced p38 signaling. We additionally examined the time-dependency of LRRK2 activation in iMicroglia (Fig. S6F-J). Following 15 minutes of exposure to 1mM LLOMe, significantly increased phosphorylation of Ser935-LRRK2, Rab10, NF-κB, and p38 was measured in treated iMicroglia relative to control cells indicating that robust cell and LRRK2 kinase responses were rapidly induced. Total LRRK2 and Iba1 levels were unaffected. The expected induction of autophagy (*25*) was demonstrated by a robust increase in LC3 lipidation following LLOMe treatment (Fig. S7A-D). LRRK2 response to LLOMe treatment was confirmed to be dependent on LRRK2 kinase activity as pre-treatment with the potent and specific kinase inhibitor MLi2 abolished increases in phosphorylation of both Ser935-LRRK2 and Rab10 (Fig. S7E-I). Collectively, these results show that iMicroglia both transcriptionally resemble brain microglia and respond to lysosomal damage with quantifiable changes in LRRK2 activity.

### The PD risk haplotype is associated with elevated LRRK2 protein expression and activity

Twelve PPMI iPSC lines were successfully differentiated to iMicroglia in six pairs of homozygous protective genotype (rs76904798-CC) or heterozygous protective:risk genotype (rs76904798-CT). LRRK2 mRNA and protein expression levels were quantified in cells at the conclusion of each differentiation experiment and the data from all six differentiation batches were aggregated. Phosphorylation of Ser935, Rab10, NF-κB and p38 proteins was measured as an indicator of LRRK2 kinase and cell activation in response to LLOMe treatment (Fig. 2C-G and Fig. S8C-D). Iba1 protein and *AIF1*, *P2RY12*, and *TMEM119* mRNA levels were used to evaluate efficiency of microglia differentiation (Fig. S8B,E-G). Significantly higher LRRK2 expression was observed in risk variant-carrying CT iMicroglia at both the mRNA and protein levels (*p*(genotype)mRNA = 2.69 ×10^-12^, *N* = 6 cell lines, *n =* 17-18 per genotype-treatment group with technical replicates, Fig. 2G; *p*(genotype)protein = 4.09 ×10^-14^, *N* = 6 cell lines, *n* = 18, Fig. 2D). We find that the rs76904798-T allele is associated with increased LRRK2 expression in iPSC-derived microglia, confirming the results of snRNAseq in the human brain.

Phosphorylation of Ser935 and of Rab10 was also significantly higher in CT cells compared to CC cells, with LLOMe treatment significantly enhancing these phosphorylation events in iMicroglia of both genotypes (*p*(genotype)S935 = 1.10×10^-4^ and *p*(treatment)S935 = 5.00×10^-24^, *N* = 6 cell lines, *n =* 18, Fig. 2E; *p*(genotype)pRab10 = 8.12×10^-5^ and *p*(treatment)pRab10 = 2.29×10^-24^, *N* = 6 cell lines, *n* = 18, Fig. 2F). LLOMe-treated iMicroglia had significantly upregulated NF-κB and p38 signaling compared to DMSO-treated control cells as expected (*p*(treatment)pNF-κB = 2.10×10^-10^, *N* = 6 cell lines, *n =* 18, Fig. S8C; *p*(treatment)p-p38 = 1.34×10^-19^, *N* = 6 cell lines, *n* = 17-18, Fig. S8D). Additionally, a significant interaction between genotype and treatment was observed for both phosphorylation of Rab10 (*p*(interaction)pRab10 = 0.009) and phosphorylation of p38 (*p*(interaction)p-p38 = 0.001), demonstrating that the impact of LLOMe treatment on iMicroglia is modulated by PD risk genotype. Taken together, these data extend the human brain data to show that microglia carrying the PD risk variant rs76904798-T not only have more expression but also more functional LRRK2, *i.e.*, that the eQTL propagates through to cellular activity.

### The LRRK2 gene is differentially accessible in microglia compared to other cell types

To begin to understand the mechanism by which LRRK2 expression is influenced by genetic variation captured by rs76904798 in microglia, we completed single nuclei ATAC sequencing (snATACseq) to study chromatin accessibility in human brain cell types. A total of 12,779 nuclei were captured from 4 frontal cortex samples, two from rs76904798-CC donors and two from rs76904798-TT donors, with an average sequencing depth of 95,000 read pairs per nucleus. The average transcription start site enrichment score was 5.3 ± 0.4 (SEM), which is acceptable by ENCODE data quality standards (*26, 27*). There were no significant differences between groups delineated by rs76904798 genotype in the number of cells measured, mean read pairs per cell, median number of fragments mapped per cell, proportion of fragments overlapping target regions, or in the proportion of transposition events in called peaks associated with cell barcodes (Fig. S9A-D). Sequenced reads in all four datasets exhibited the expected fragment size distribution, with periodicity of peaks corresponding to nucleosome-free regions (<100 bp), mononucleosome (∼200 bp), and dinucleosome (∼400 bp) bound fragments (*28*) (Fig. S9E). There was also evidence of TSS enrichment in all datasets (Fig. S9F). After exclusion of outlying nuclei, the four datasets were merged into a single dataset containing 7,805 high quality nuclei. The total number and proportion of fragments encompassing peaks, the proportion of fragments mapping to blacklisted regions, and the nucleosome signal for the nuclei included in the merged dataset are displayed in Fig. S9G-J.

Sixteen distinct cell populations were identified by shared nearest neighbor modularity optimization clustering analysis. Nuclei originating from all four donors contributed to each cluster, with the exception of cluster 5 that did not contain any nuclei derived from one donor. Clusters were not distinguished by donor genotype at rs76904798. Differential chromatin accessibility at marker gene loci was used to assign a cell type to each cluster. Clusters 14 and 15 were excluded from downstream analysis due to increased accessibility near marker genes of multiple cell types, resulting in 14 defined cell populations composed of 7,732 nuclei in total (Fig. 3A-B). Cell type assignments were validated by integration of the merged snATACseq dataset with a merged snRNAseq dataset containing nuclear transcriptomes from 16,148 frontal cortex nuclei extracted from the same four donors. Clustering analysis and cell population identification were completed for this subset (*N* = 4) of the full snRNAseq dataset (*N* = 15). Using the frontal cortex snRNAseq subset as a reference, nuclei in the snATACseq dataset were assigned a predicted cell type dependent on pair-wise comparison of gene expression and gene accessibility between two nuclei in each respective dataset (Fig. S10A-B). The predicted labels matched the cell type assignments made based on marker gene accessibility for 99.94% of nuclei (7,727 out of 7,732), with one apparently incorrect label of ExN.6 assigned to five nuclei within an inhibitory neuron population. The overall agreement between the cell types predicted by snRNAseq integration and those assigned based on differential accessibility of marker genes indicates that our cell type assignments are robust.

**Fig. 3.**
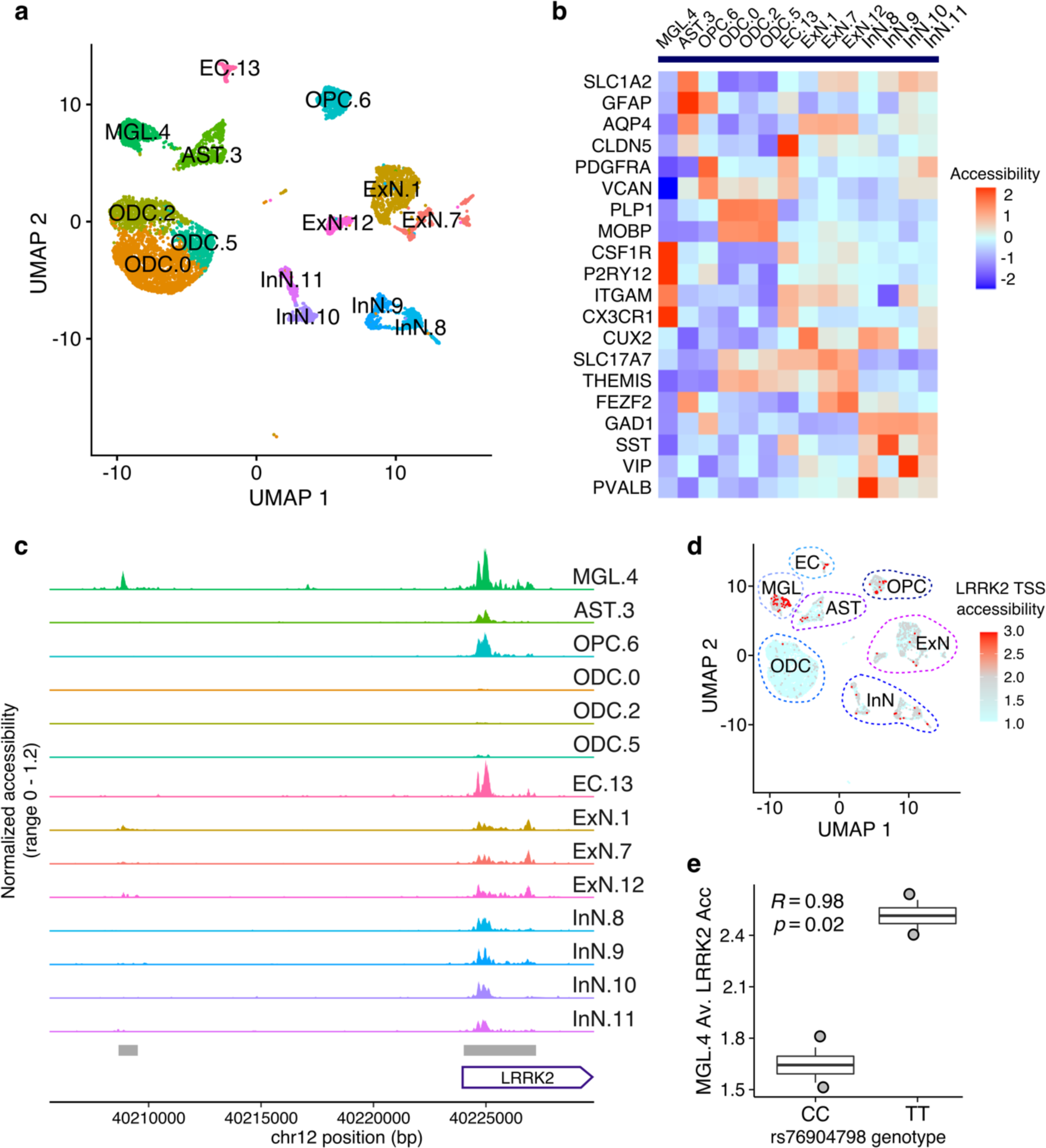
Chromatin in the *LRRK2* region is differentially accessible in brain microglia. (A) UMAP visualization of cell populations identified in four human frontal cortex samples by snATACseq. (B) Heatmap indicating the relative chromatin accessibility near known cell type marker genes in each cell population. (C) Chromatin landscape of the *LRRK2* promoter region representing average accessibility of cells within each cluster. Significantly accessible regions, or called peaks, are indicated by gray bars. (D) UMAP plot showing significantly increased accessibility of the *LRRK2* transcription start site (TSS) region, chr12:40,224,010-40,227,222 in microglia population “MGL.4” compared to all other cell types (adjusted *p* = 1.57 x 10^-57^, average log2(fold change) = 0.89). (E) Boxplot where each point represents average *LRRK2* accessibility of cells in “MGL.4” contributed by each of the four donors. Using the number of minor alleles (rs76904798-CC = 0, rs76904798-TT = 2) as the x-axis in a linear regression model, Pearson’s *R* = 0.976 (*p* = 0.024, df = 2, *t* = 6.303).

Having established that all expected brain cell types were detected by snATACseq, chromatin accessibility around the *LRRK2* gene in each cell type was examined. Pseudo-bulk tracks representing average chromatin accessibility within each cluster in the *LRRK2* promoter region are shown in Fig. 3C. The called peak in the *LRRK2* transcription start site region (coordinates chr12:40,224,010-40,227,222) is significantly more accessible in the microglia population MGL.4 compared to all other brain cell types (adjusted *p* = 1.57×10^-57^, likelihood ratio test comparing a logistic regression model to a null model followed by Bonferroni multiple test correction; *N* = 4 frontal cortex samples; *n*MGL.4 = 521, *n*AST.3 = 531, *n*EC.13 = 142, *n*OPC.6 = 479, *n*ODC.0 = 1772, *n*ODC.2 = 842, *n*ODC.5 = 503, *n*ExN.1 = 1107, *n*ExN.7 = 400, *n*ExN.12 = 228, *n*InN.8 = 361, *n*InN.9 = 327, *n*InN.10 = 275, *n*InN.11 = 244 nuclei; Fig. 3D). Interestingly, the average chromatin accessibility in and around the *LRRK2* gene was significantly higher in rs76904798-TT cells relative to rs76904798-CC cells in population MGL.4 (*p* = 0.025, by two-tailed unpaired t-test with Welch’s correction, t = −6.303, df = 1.97, α = 0.05). Linear regression analysis of the average *LRRK2* gene accessibility in nuclei contributed by each donor indicated that accessibility was significantly correlated with presence of the T allele (Pearson’s *R* = 0.98, *p* = 0.024, *N* = 4, t = 6.303, df = 2, Fig. 3E). Though a trend towards increased *LRRK2* gene accessibility in TT cells was observed in other cell populations, the difference was statistically significant by either measure only in microglia population MGL.4 (Fig. S10C-D). The increased number of fragments mapping to the *LRRK2* gene body in TT cells compared to CC cells is visualized in Fig. S10E. These results indicate that the T allele at rs76904798 is correlated with increased accessibility throughout the *LRRK2* gene body in microglia. Thus, the likely mechanism by which the eQTL is restricted to microglia is via chromatin regions that are most accessible in microglia.

### Regions of differentially accessible chromatin in brain microglia are reproduced in iMicroglia

To determine how faithfully the iMicroglia cell model reproduces the DNA architecture observed in brain microglia, we performed bulk ATACseq using iMicroglia differentiated from PPMI cell lines, two of genotype rs76904798-CC and two of genotype rs76904798-CT. We also performed bulk ATACseq of iPSC-derived forebrain neurons (iFbN) for comparison. In parallel, the snATACseq dataset was split into separate aligned fragment files for each identified cell population. We then completed an integrated differential accessibility analysis using called peak counts from each of these datasets (Fig. 4A). We found that genome-wide chromatin accessibility was highly similar between iMicroglia and brain microglia (Pearson’s *R* = 0.82-0.86 in different cell lines) and that the iMicroglia clustered with the brain microglia using complete linkage clustering (Fig. 4B). The iFbN equally clustered near the brain inhibitory neuron populations (Pearson’s *R* = 0.82-0.85). The same peaks of chromatin accessibility detected in brain microglia in the *LRRK2* region were captured in the iPSC-derived microglia but were absent in the iFbN (Fig. 4C). These data demonstrate that iMicroglia faithfully replicate the microglia-specific differential chromatin accessibility in the *LRRK2* region.

**Fig. 4.**
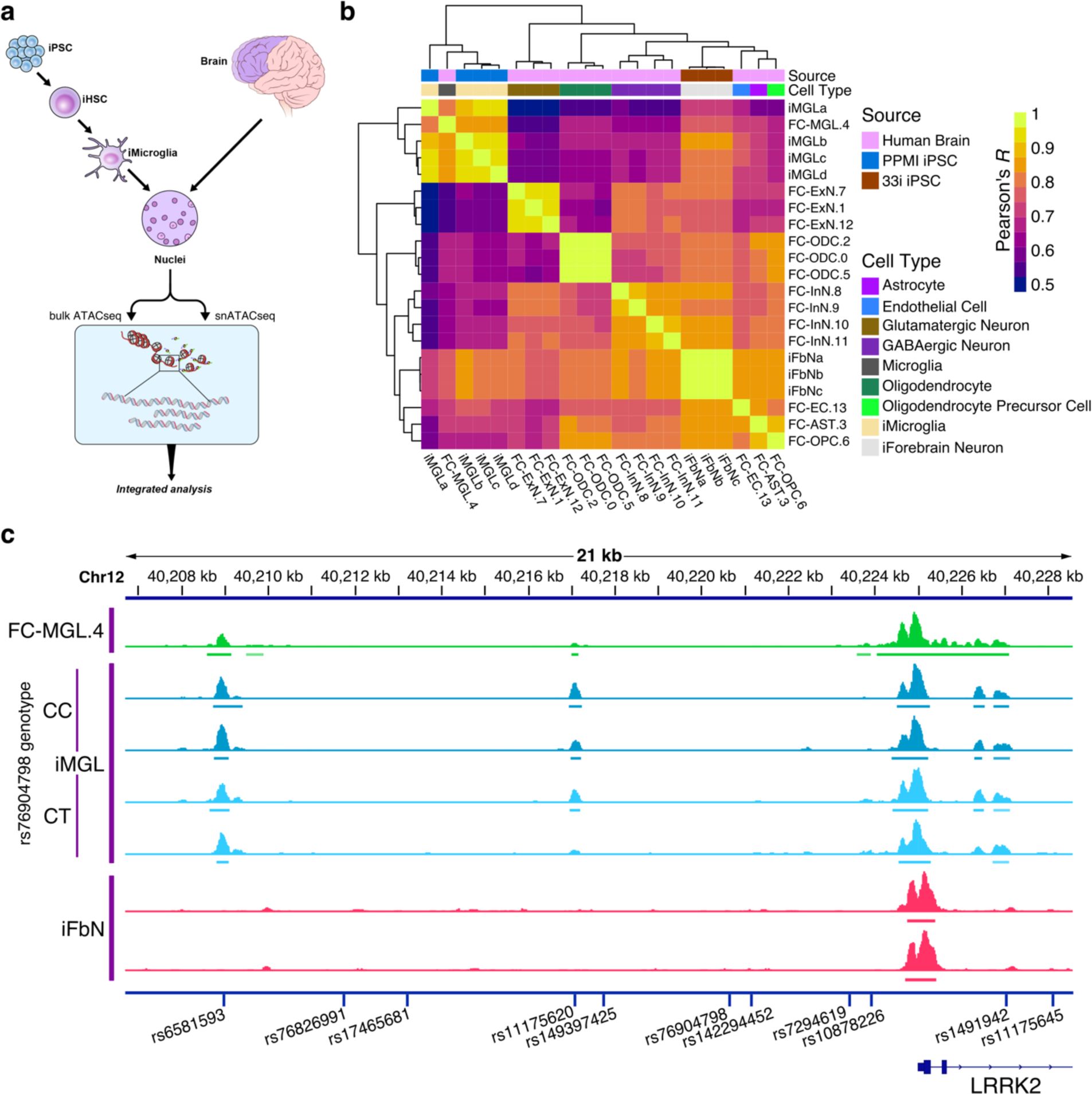
Chromatin architecture of iMicroglia resembles that of brain microglia in the *LRRK2* region. (A) Diagram illustrating the integrated analysis of bulk ATACseq of iMicroglia with snATACseq of human frontal cortex. Graphic design credit to Erina He at NIH Medical Arts. (B) Heatmap showing the results of DESeq2 analysis of peak counts for iPSC-derived or brain cell types, with hierarchical clustering of Pearson correlation values using the complete-linkage method. For iMGLa vs FC-MGL.4, Pearson’s *R* = 0.820, df = 164,188, *t* = 579.49, *p* = 2.22 x 10^-16^. (C) Comparison of the chromatin landscape of the *LRRK2* promoter region in brain microglia (population “MGL.4”) by snATACseq vs. that observed in iMicroglia via bulk ATACseq of four differentiated iPSC lines (PPMI 3411, 3453, 3446, and 4101) and iFbn via bulk ATACseq of one differentiated iPSC line (33i, with three technical replicates). ATACseq alignments were visualized using the Broad Institute’s Integrative Genomics Viewer^58^. Called peaks are represented by horizontal bars under each track. Locations of SNPs in linkage disequilibrium with PD-risk SNP rs76904798 are shown.

### The rs76904798 variant does not drive the observed LRRK2 eQTL in iMicroglia

To test the hypothesis that the rs76904798 variant is responsible for the observed microglia-specific LRRK2 eQTL, we used CRISPR/Cas technology to exchange the major/protective variant (C) for the minor/risk variant (T) in an iPSC line homozygous for the reference allele. Guide RNA (gRNA) and donor template design is illustrated in Fig. S11A, with confirmation of edited clone genotypes by targeted amplicon sequencing. Correctly edited cell lines containing only the target single nucleotide substitution, and not other changes in the immediate (300bp) vicinity, were differentiated to iMicroglia alongside the original parental cell line and a “non-edited” control (NEC) cell line.

Assays of LRRK2 expression level in edited iMicroglia indicated that LRRK2 expression was unaffected by introduction of the risk variant at rs76904798 (Fig. S11B-F). LRRK2 expression was similar between edited iMicroglia and parental iMicroglia at both mRNA and protein levels. Likewise, LRRK2 response to LLOMe treatment was similar between edited cells and non-edited cells (Fig. S12) indicating that absence of an effect was not a consequence of the LRRK2 assays having an upper limit of saturation. These data demonstrate that the lead GWAS variant, rs76904798[C/T], itself does not underlie the microglia-specific rs76904798:LRRK2 eQTL.

## Discussion

Here, we have shown that expression as a quantitative trait in a locus associated with human PD is mediated through microglia and microglial-specific epigenetic events. Importantly, the cell type specificity of the eQTL differs from baseline patterns of expression, which are more prominent in excitatory projection neurons and oligodendrocyte precursors. If generalized, this data would indicate that cell type-specific eQTL mapping may be critical in resolving GWAS signals where common genetic variants have been associated with a range of phenotypes.

A critical assumption of the current analysis is that at this locus we can confidently infer the identity of the causal gene, *LRRK2*. This inference rests on the observation that coding variants in *LRRK2* are causal for late onset PD in families, albeit with age-dependent and incomplete penetrance. It has been previously proposed that such pleomorphic risk loci (*29*) represent differing genetic variants that drive disease by quantitatively different extents but in the same direction. In the case of *LRRK2*, the overwhelming evidence indicates that coding variants induce a gain of function of activity that is associated with disease. Our observations that microglia from human donors with higher GWAS-nominated disease risk have higher net expression and activity of LRRK2, in combination with prior data from monocytes and derived cells ex vivo (*9*), indicate that a common mechanism of all variants at the *LRRK2* locus is to enhance function. In this context, it is worth noting that analyses using the human brain have not reported a strong eQTL but do identify a signal in blood. We infer that there is a shared eQTL between microglia and macrophages at the *LRRK2* locus that would likely be difficult to detect in the brain where many other cell types are present that express LRRK2 but do not propagate the eQTL.

It is equally of interest that we did not find evidence of an eQTL in excitatory neurons or oligodendrocyte precursor cells, despite levels of expression that are higher than in microglia. We infer that there are at least two independent mechanisms that control LRRK2 expression, resulting in both genotype-independent and -dependent expression. The observation that there are differential regions of open chromatin between microglia and neurons provides a potential to mechanistically resolve these differences. Our assumption is that the open chromatin regions in microglial DNA allow for specific regulation of *LRRK2* expression and that the differences between basal and regulated expression in different cell types may be biologically important. By extension, this suggests that there may be subtleties in function of this molecule in different cell types in the human brain, although this hypothesis will require further experimental validation. However, we do not claim that other cell types are not involved in the pathology of human *LRRK2* associated disease, and note that non-cell autonomous events are likely mediators of a multi-brain region, age-related phenotype.

We were able to replicate the key observations of a microglial eQTL at *LRRK2* in human iPSC-derived cells, which we were able to demonstrate have a transcription program and regions of open chromatin similar to microglia in the human brain. This then allowed us to evaluate two critical events that would not currently be feasible to address in a non-experimental system such as the human brain. First, we could show that the increase in mRNA expression encoded by the microglial eQTL propagates through the level of protein expression and activity. Although enzyme activity as a quantitative trait would be predicted to be an emergent property of genotype-dependent mRNA expression, experimental and quantitative verification in a restricted cell type excludes that the eQTL is counteracted by homeostatic regulation of protein and activity in cells. Second, we were able to control the activation state of cells, which again would be impossible to perform in the human brain. Here, we extend our recent observation that lysosomal damage can activate the LRRK2 signaling pathway (*24*) to show that activation can occur in iMicroglia and that activation shows interaction with genotype. Given results from other laboratories that inflammatory signaling can promote LRRK2 activation through a similar pathway (*30*), we infer that neuroinflammation may interact with genotype at *LRRK2*, and potentially at other lysosomal genes associated with PD (*31*), to influence overall brain phenotype. Such interactions may be further modified by aging, which influences both inflammation and protein degradation pathways (*32*). Thus, net lifetime risk of a sporadic disease such as PD may emerge from a combination of genetic risk factors, aging, and non-genetic events distributed across diverse cell types.

We also found that the most significantly associated SNP in PD GWAS did not fully explain the microglial eQTL, finding no difference using CRISPR/Cas editing from protective to risk variant in iPS cells subsequently differentiated to iMicroglia. It is likely that the lead or sentinel SNP at this and other GWAS loci captures a complex set of linked variants that may each contribute to differential expression. Such haplotypic effects may prove complicated to resolve using single editing approaches, and might require replacement of large genomic regions. In addition, while the current analysis focuses on a relatively common variant that can be reliably evaluated by GWAS, it remains possible that rare variants on a common haplotype may represent true causal alleles. Thus, iterative analyses between better powered GWAS and experimental systems are likely required to further resolve disease associations to the single base pair level.

Overall, we present an integrated analysis of a genomic locus associated with PD in humans and identify microglia as important mediators of genetic risk via a cell type-enhanced eQTL. Whether such events can be shown to generalize across loci and other diseases remains to be established, but these data suggest that baseline expression on a per cell type basis does not predict contributions to eQTL. Further work is therefore needed to resolve single cell and single variant mediators in complex diseases.

## Acknowledgments

This work utilized the computational resources of the NIH HPC Biowulf cluster (http://hpc.nih.gov). Human tissue was obtained from the NIH NeuroBioBank. We are grateful for the services and the expert advice of the NIH NHLBI Flow Cytometry Core, the NIH NIDDK Genomics Core, the NIH NCI Single Cell Analysis Facility, the NISC Comparative Sequencing Center, and Erina He at the NIH Medical Arts Branch. We thank Dr. Steve Finkbeiner (Gladstone Institutes) for the gift of the CS83iCTR-33n1 iPSC line and Dr. Changyoun Kim (NIH) for the gift of α-synuclein conditioned medium.

## Funding

This research was supported in part by the Intramural Research Program of the NIH, National Institute on Aging, and by the Michael J. Fox Foundation. PPMI is funded by The Michael J. Fox Foundation for Parkinson’s Research and corporate sponsors (https://www.ppmi-info.org/about-ppmi/who-we-are/study-sponsors/).

## Author contributions

R.G.L. designed, performed, and analyzed experiments. A.B. and X.R. advised on and performed experiments. A.B.S., C.B. and J.R.G. advised on data analysis and interpretation; M.R.C advised on design and interpretation of experiments. R.G.L. and M.R.C. wrote the manuscript. A.B., X.R., C.B., and J.R.G. edited the manuscript.

## Data and materials availability

Human frontal cortex snRNAseq and snATACseq, iMicroglia scRNAseq, and iMicroglia and iFbN bulk ATACseq datasets have been deposited in the Gene Expression Omnibus (GEO) database under accession number GSE163323. Code for data analysis and visualization is available in the github repository https://github.com/neurogenetics/LRRK2-microglia-eQTL/.

## Materials and Methods

### Acquisition of human brain tissue

Frozen post-mortem human frontal cortex samples from donors without known neurological disease were obtained from the University of Maryland Brain and Tissue Bank through the NIH NeuroBioBank. Genome-wide genotyping and genome sequencing of these brain tissues was completed previously by the North American Brain Expression Consortium (dbGaP Study Accession: phs001300.v1.p1) (*33*). Five donors of each genotype at rs76904798[CC/CT/TT] were selected to yield a sample group balanced across age, sex, and genotype (Male:Female = 8:7, age range = 16-61 years old, median age = 39, CC:CT:TT = 1:1:1). Donor demographic data are presented in Table S1.

### Nuclei isolation from frozen human brain

Nuclei extraction from the fifteen frontal cortex samples was completed in four batches that were similar in terms of sample count (n = 3 or 4), average number of risk alleles (1), average age (35.3 ± 3.1 years), average post-mortem interval (16.1 ± 1.9 hours), and average sex (1.5 ± 0.1, where Male was coded as 2 and Female as 1). For each sample, approximately 100mg of tissue was thawed in 2mL of cold lysis buffer (Nuclei PURE Lysis Buffer/1mM DTT/0.1% Triton X-100, Sigma Aldrich #NUC201) and homogenized using a Dounce homogenizer. The homogenate was vortexed briefly with an additional 8mL of lysis buffer and incubated for 10 minutes on ice. After pipetting 10mL of cold 1.8M Sucrose Cushion Solution (Sigma Aldrich #NUC201) into an ultracentrifuge tube (Beckman Coulter #344058) on ice, tissue lysate was resuspended with 18mL of cold 1.8M Sucrose Cushion Solution and added slowly with minimal disruption of the bottom layer of solution. Samples were centrifuged for 45 minutes at 30,000 x *g* at 4 ℃ in a pre-cooled ultracentrifuge. The pelleted nuclei were resuspended in 1mL ice-cold Nuclei Suspension Buffer (NSB, PBS with 0.01% BSA, New England BioLabs #B9000S, and 0.1% SUPERase-In RNase inhibitor, ThermoFisher Scientific #AM2696) as described by Habib *et al* 2017 (*17*). The nuclei were washed twice in a total of 5mL of ice-cold NSB, with centrifugation for 5 minutes at 500 x *g* at 4 ℃. Nuclei were resuspended in 110#μL of cold NSB and counted using an automated cell counter (Bio-Rad). Concentration was adjusted to ∼1000 nuclei/#μL.

### Single nuclei RNA sequencing of human brain

The isolated nuclei were submitted to the Single Cell Analysis Facility (Center for Cancer Research, National Cancer Institute) for single nuclei RNA sequencing (snRNAseq) library construction using the Chromium Single Cell Gene Expression Solution v3 (10x Genomics) targeting 6,000 nuclei for capture. Pooled sequencing libraries were loaded at a concentration of 1.8pM with 10% PhiX spike-in and sequenced using eight Illumina NextSeq 150 Cycle Hi-Output v2.5 kits (Illumina #20024907) on an Illumina NextSeq 550 System, with a targeted sequencing depth of around 33,000 reads per nucleus. Feature-barcode matrices were generated for each sample following sequence alignment and counting by Cell Ranger software (10x Genomics).

Analyses of these data were completed using the Seurat package (version 3.1) (*34*) in R version 4.0.0 on the Biowulf cluster provided by the NIH High Performing Computation (HPC) group (http://hpc.nih.gov). SCTransform (*35*) normalization and regression of mitochondrial genes were performed on each dataset prior to integration based on pair-wise expression of “anchor” genes (*36*). Shared-nearest-neighbor-based (*37*) clustering of the integrated dataset was completed using a clustering resolution parameter of 1 and clusters were visualized following dimensional reduction via Uniform Manifold Approximation and Projection (UMAP) (*38*). Differential gene expression analysis was used to characterize the composition of each cell cluster. Clusters were assigned a cell type based on significantly differentially increased expression of the following marker genes: *SLC17A7* (excitatory neuron, ExN); *GAD1* (inhibitory neuron, InN); *CLDN5* and *COLEC12* (endothelial cell, EC); *PDGFRA* and *OLIG2* (oligodendrocyte precursor cell, OPC); *PLP1* and *MOBP* (oligodendrocyte, ODC); *SLC1A2*, *SLC1A3*, *GFAP*, and *AQP4* (astrocyte, AST); *P2RY12*, *ITGAM*, *CSF1R*, and *CX3CR1* (microglia, MGL) in accordance with known genetic signatures of human brain cell types (*11, 17–19*). Differentially expressed genes were defined as those with significantly different expression levels between two groups of cells as measured by a Wilcoxon rank sum test with Bonferroni correction. Correlation between genotype at rs76904798 and *LRRK2* expression was assessed by simple linear regression comparing average expression of *LRRK2* in nuclei of each cluster per donor with dosage of the risk allele rs76904798-T (donor genotype rs76904798-CC = 0, rs76904798-CT = 1, or rs76904798-TT = 2).

### Acquisition of induced pluripotent stem cell lines

Patient-derived induced pluripotent stem cell (iPSC) lines, reprogrammed from peripheral blood mononuclear cells, were obtained from the Golub Capital iPSC Parkinson’s Progression Markers Initiative (PPMI) Sub-study (www.ppmi-info.org/cell-lines). Genome sequencing of these lines was completed previously and is available at https://www.ppmi-info.org/. No cell lines homozygous for the risk variant at rs76904798 (genotype = TT) were available. Six iPSC lines heterozygous for the risk variant at rs76904798 (genotype = CT) and without known PD-associated mutations (*e.g.* LRRK2 G2019S) were matched with an iPSC line not carrying the risk variant (genotype = CC) based on sex and birth year of the donor (Male:Female = 1:1, average donor birth year = 1949 ± 9). Additional characteristics of these cell lines are summarized in Table S2. The CS83iCTR-33n1 (33i) episomal iPSC line, generated from fibroblasts at Cedars-Sinai, was kindly shared by Steve Finkbeiner (Gladstone Institutes). Commercial human iPSC line A18945 was obtained from ThermoFisher (#A18945). A LRRK2-knockout version of this iPSC line (A18945 LRRK2 KO) was generated previously using the CRISPR-Cas9 system. Two gRNAs (5’-GGCCTATGAAGGAGAAGAAG-3’ and 5’-GATTAATTGCTGCAATAAGG-3’) targeting LRRK2 sequence near GTP binding and ATP kinase binding domains were synthesized and cloned into pSpCas9(BB)-2A-GFP plasmid (Addgene #48138). Plasmids were transfected into A18945 iPSC line using Lipofectamine-Stem transfection reagent (ThermoFisher #STEM00001) and GFP-positive cells were collected by fluorescence-activated cell sorting (FACS) after 24 hours. Sorted cells were further cloned into single cells, then expanded and assessed by Western blot for LRRK2 expression.

### Differentiation of iPSC to iMicroglia

iPSC lines were differentiated to iMicroglia in pairs. Prior to differentiation, cells were cultured in TeSR-E8 medium (StemCell Technologies #05990) in matrigel-coated (Corning #354230) 6-well plates. iPSC were first differentiated to hematopoietic stem cells (iHSC) using the STEMdiff Hematopoietic Kit from StemCell Technologies (#05310) incorporating protocol adaptations recommended by the Blurton-Jones lab (*21*). Briefly, culture medium was replaced with 2 mL/well iHSC Medium A (StemCell Technologies #05310) on differentiation Day 0. An additional 1 mL/well of iHSC Medium A was added on Day 2. On Day 3, medium was replaced with 2 mL/well iHSC Medium B (StemCell Technologies #05310). An additional 1 mL/well of iHSC Medium B was added every other day. On Day 11 or 12, iHSC were collected using a serological pipette and transferred to a 50 mL conical tube along with a rinse of 1 mL DPBS (ThermoFisher #14190250) per well. After centrifugation at 300 x *g* for 5 minutes, iHSC were re-plated in matrigel-coated 6-well plates at a density of up to 600,000 cells per well for iMicroglia differentiation according to the protocol published by the Blurton-Jones lab (*21, 22*).

Cells were fed every other day for 24 days with iMicroglia Differentiation Medium consisting of iMicroglia Basal Medium (1% (v/v) nonessential amino acids, ThermoFisher #11140050; 1% (v/v) GlutaMAX, ThermoFisher #35050061; 0.1% (v/v) 5 mg/mL human insulin, Sigma Aldrich #I2643-50mg; 2% (v/v) B27, ThermoFisher #17504044; 2% (v/v) ITS-G, ThermoFisher #41400045; 0.5% (v/v) N2, ThermoFisher #17502048; 0.04% (v/v) 1M monothioglycerol, Sigma Aldrich #M1753-100ML; DMEM/F-12 without (–) phenol red, ThermoFisher #11039021) plus 100 ng/mL IL-34 (PeproTech #200-34), 50 ng/mL TGFβ1 (PeproTech #100-21), and 25 ng/mL M-CSF (PeproTech #300-25). On iMicroglia differentiation day 25, cells were collected and centrifuged at 300 x *g* for 5 minutes, then resuspended in 1 mL/well of iMicroglia Maturation Medium made up of iMicroglia Differentiation Medium plus 100 ng/mL CD200 (Novoprotein #C311) and 100 ng/mL CX3CL1 (PeproTech #300-31). An additional 1 mL/well of iMicroglia Maturation Medium was added every other day until iMicroglia (iMGL) were collected for experiments, as early as day 28.

Preliminary experiments were completed with A18945 WT and LRRK2 KO iPSC lines to confirm expression of microglia markers and LRRK2 in differentiated cells. Floating cells were collected in culture medium then wells were rinsed with 1 mL DPBS which was added to the same collection tube. The remaining adherent cells were incubated with Accutase (StemCell Technologies #07920) for 3 to 5 minutes at 37°C and collected into a separate 15mL tube containing 5mL of DMEM/F-12 (–) phenol red. Cells were pelleted by centrifugation at 300 x *g* for 5 minutes and submitted to western blot analysis. Floating and adherent iMicroglia populations were combined for all later experiments.

### Differentiation of iPSC to forebrain neurons

To generate induced forebrain neurons (iFbN) CS83iCTR-33n1 (33i) iPSC were grown in E8 medium (ThermoFisher #A1517001) on matrigel-coated plates. Once confluent, cells were differentiated as previously published (*39*). Briefly, media was changed to neuronal differentiation media N3 (50% DMEM/F-12 (ThermoFisher #11320-033), 50% Neurobasal (ThermoFisher 21103-049) with 1x Pen Strep (ThermoFisher #15140-122), 1x B-27 minus Vitamin A (ThermoFisher #12587010), 1x N-2 Supplement (ThermoFisher #17502048), 1x GlutaMAX (ThermoFisher, #35050061), 0.5x MEM Nonessential Amino Acids (ThermoFisher, #11140050), 55uM 2-mercaptoethanol (ThermoFisher #21985023) and 1ug/ml Insulin (Millipore Sigma #91077C-250mg)). N3 was changed daily and supplemented with 1.5uM Dorsomorphin dihydrochloride (Tocris Bioscience #3093) and SB431542 (Stemgent #04-0010-05) for days 1 through 11. Days 12 through 15 received N3 without additional supplements and beginning on day 16 0.05uM Retinoic acid (Millipore Sigma #R2625) was added to the basal N3 media. Cells were split 1:2 and plated on 0.01% Poly-L-ornithine (Millipore Sigma #P4957), 2ug/mL Fibronectin (Fisher Scientific #CB40008A) and 0.2ug/mL Laminin (Millipore Sigma #L6274) coated plates in N4 media (N3 with 0.05uM Retinoic acid, 2ng/ml BDNF (R&D Systems #248-BDB) and 2 ng/ul GDNF (R&D Systems #212-GD)). iFbN were replated on day 24 in a coated 96 well plate at 75000 cells/well. N4 media was changed every 2 days until collected for experiments.

### Assays of mRNA and protein expression levels

For quantitative PCR (qPCR) experiments, iMicroglia were collected by centrifugation at 6000 x *g* for 1.5 minutes. RNA was extracted from pelleted cells using TRIzol reagent (ThermoFisher #15596018). cDNA was prepared using the SuperScript III First-Strand Synthesis SuperMix for qRT-PCR (ThermoFisher #11752-050). FAM-MGB-labeled TaqMan Gene Expression Assays for *LRRK2*, *AIF1*, *P2RY12*, and *TMEM119* (#4331182: Hs01115057_m1, Hs00610419_g1, Hs01881698_s1, and Hs01938722_u1) along with VIC-MGB-labeled *PPID* (#4448489: Hs00234593_m1) were obtained from ThermoFisher. Samples were run in quadruplicate using TaqMan Fast Advanced Master Mix (ThermoFisher #4444557) on a QuantStudio 6 Flex Real-time PCR system (Applied Biosystems) in 384-well plates. mRNA levels were calculated by normalization to those of housekeeping gene cyclophilin D (*PPID*) as previously described (*40*).

For immunoblotting, pelleted cells were resuspended in 1x Cell Lysis Buffer (Cell Signaling #9803) containing 1x Halt phosphatase and protease inhibitor cocktails (ThermoFisher #78426 and #78429) and incubated on ice for 20-30 minutes. Lysates were centrifuged at 16,000 x *g* for 10 minutes, and cleared lysates were separated by SDS/PAGE on 4-20% Criterion TGX precast gels (BioRad #5671095). Immunoblots were probed overnight with the following primary antibodies at a 1:2000 dilution: anti-LRRK2 (Abcam #ab133474), anti-pS935-LRRK2 (Abcam #ab133450), anti-NF-κB (CellSignaling #8242), anti-phospho-NF-κB (CellSignaling #3033), anti-p38 (CellSignaling #9212), anti-phospho-p38 (CellSignaling #9211), anti-Rab10 (AbCam #ab237703), anti-phospho-Rab10 (AbCam #ab230261), anti-Iba1 (Wako 016-20001), anti-P2Y12 (ThermoFisher #702516), anti-LC3B (CellSignaling #2775), anti-GAPDH (Sigma #G9545), anti-actin (Sigma #A3853), and anti-cyclophilin b (AbCam #ab16045). Both mRNA and protein expression levels were compared between genotype and treatment groups by one-way or two-way analyses of variance (ANOVA) with Tukey’s, Dunnett’s, or Sidak’s multiple comparisons tests. Statistical tests were performed in R version 4.0.0 or in Prism version 7.0d.

### Single cell RNAseq of iMicroglia

iMicroglia differentiated from two PPMI cell lines were submitted to single cell RNA seq (scRNAseq) analysis on iMicroglia differentiation day 28. Floating cells were collected in culture medium, then adherent cells were collected into the same tube following a 3- to 5-minute incubation at 37 °C with Accutase. After centrifugation at 300 x *g* for 5 minutes, cell pellets were resuspended in 1 mL FACS buffer (DPBS with 2% BSA, Miltenyi Biotec #130-091-376, plus 0.05mM EDTA, ThermoFisher #15575020, and 5 ng/mL SCF, ThermoFisher #PHC2115) and incubated with Zombie Violet Fixable Viability dye (BioLegend #423113) at a 1:1000 dilution for 5 minutes at room temperature. Stained cells were washed with 10mL DPBS, centrifuged for 5 minutes at 300 x *g*, and resuspended in 500 μL FACS buffer. After filtering through a 35 μm mesh tube cap (Corning #352235), purification of live cells by FACS was completed by the NHLBI Flow Cytometry Core.

The collected live iMicroglia were pelleted by centrifugation at 300 x *g* for 5 minutes, resuspended in 500 μL PBS/0.01% BSA/0.1% SUPERase-In RNase Inhibitor, and counted using a LUNA-FL automated fluorescent cell counter (Logos Biosystems). The cell suspension was diluted to a concentration of 1000 cells/μL with additional PBS/0.01% BSA/0.1% SUPERase-In RNase Inhibitor. The Chromium Single Cell Gene Expression Solution v3.1 (10x Genomics) was used to construct scRNAseq libraries for each sample targeting 5,000 nuclei for capture.

Sequencing of the pooled libraries was completed by the NIDDK Genomics Core on an Illumina NextSeq 550 system using a NextSeq 150 Cycle Hi-Output v2.5 kit (Illumina #20024907), generating a total of 400 million reads for an estimated sequencing depth of 40,000 reads per cell. Raw sequencing data were aligned and counted using Cell Ranger software (10x Genomics). Normalization, integration, and clustering analysis of these two scRNAseq datasets was completed using the Seurat package (version 3.1) (*34*) in R as previously described. *iMicroglia treatment assays*

Floating and adherent iMicroglia were collected on differentiation day 28-29 and replated 24-48 hours before treatment in uncoated ultra-low attachment 24-well plates (Corning, Sigma #CLS3473-24EA) in 400 μL iMicroglia Experiment Medium (iMicroglia Maturation Medium without TGFβ1) at a density of 1-3 x 10^5^ cells per well. Treatment mediums were prepared at 5x concentration in DMEM/F12 (–) phenol red. These included 500 ng/mL lipopolysaccharides from *Escherichia coli* (LPS, Sigma Aldrich #L4391), 5x α-synuclein conditioned medium (*41*) (aSCM) diluted from a 618x stock generously shared by Changyoun Kim (Laboratory of Neurogenetics, National Institute on Aging), and 5mM of lysosomal damaging compound and LRRK2 activator (*24*) L-leucyl-L-leucine methyl ester (LLOMe, Sigma Aldrich #L7393) dissolved in DMSO. 100μL of 5x medium was added to iMicroglia in their existing culture medium resulting in 100 ng/mL LPS, 1x aSCM, and 1mM LLOMe treatment concentrations. Treated cells were incubated at 37 °C for the described times before being collected for western blot analysis.

Comparison of treatment conditions was completed using iMicroglia differentiated from two PPMI cell lines, one WT-LRRK2 (PPMI 3448) and one carrying the kinase activating G2019S-LRRK2 mutation (PPMI 51782) to serve as a positive control for detection of LRRK2-mediated phosphorylation events. A LLOMe time course assay was completed using iMicroglia differentiated from a G2019S-carrying iPSC line (PPMI 51440) to determine the optimal treatment time. To assess dependency of LLOMe treatment response on LRRK2 kinase activity, iMicroglia differentiated from three PPMI cell lines were subjected to a 2-hour pre-treatment with selective LRRK2 kinase inhibitor MLi-2 (*42, 43*) prior to LLOMe treatment. MLi-2 (Tocris #5756) was reconstituted in DMSO to a stock concentration of 10mM. 100 μL of 5μM MLi-2 in DMEM/F12 (–) phenol red were added to iMicroglia re-plated in 400 μL of iMicroglia Experiment Medium to achieve a final MLi-2 treatment concentration of 1μM. After incubating for 2 hours at 37 °C, 100 μL of 6mM LLOMe treatment medium were added to the pre-treated wells, resulting in a treatment concentration of 1mM LLOMe, and cells were incubated for 15 minutes before collection for analysis. A volume of DMSO equal to that of the applied MLi-2 and/or LLOMe stock solutions in DMEM/F12 (–) phenol red was added to untreated wells of cells as a negative control. Cells were collected for western blot analysis following treatment. iMicroglia differentiated from the six pairs of PPMI cell lines listed in Table S2 were submitted to a 15 minute treatment with 1mM LLOMe prior to collection for western blot and qPCR analysis.

### Single Nuclei ATAC sequencing of human brain

Single nuclei ATAC sequencing (snATACseq) was performed on frontal cortex samples from 4 of the 15 donors included in the snRNAseq experiment, two with genotype rs76904798-CC and two with genotype rs76904798-TT (M:F = 1:1 in each genotype group). Donors from whom samples were used for both snRNAseq and snATACseq are highlighted in Table S1. Nuclei were isolated from frozen tissue as described for snRNAseq. Following ultracentrifugation, pelleted nuclei were resuspended in 1 mL ice-cold Nuclei Wash Buffer (NWB) made up of 10mM Tris-HCl pH 7.5 (KD Medical #RGF-3350), 10mM NaCl (KD Medical #RGF-3270), 3mM MgCl2 (ThermoFisher AM9530G), 1% BSA (Miltenyi Biotec #130-091-376) and 0.1% Tween-20 (Sigma Aldrich #P7949) in nuclease-free water as recommended by 10x Genomics (*44*).

Suspended nuclei were washed twice with 5 mL of ice-cold NWB, with 5-minute centrifugation at 500 x *g* at 4 ℃. Nuclei were resuspended in 100#μL of cold Diluted Nuclei Buffer (10x Genomics) and counted using a LUNA-FL automated fluorescent cell counter (Logos Biosystems). The nuclei suspension was further diluted to a concentration of 2500 nuclei/#μL.

Sequencing libraries were constructed using the 10x Genomics Chromium Single Cell ATAC Solution v1.1 targeting 6,000 nuclei for capture. Illumina-compatible libraries were submitted to the NIH Intramural Sequencing Center (NISC) for sequencing. Libraries were pooled based on qPCR quantification, then sequenced on a NovaSeq 6000 DNA Sequencer (Illumina) on an SP flow cell with a loading concentration of 300pM to achieve a targeted sequencing depth of at least 25,000 read pairs per nucleus. The resulting FASTQ files were aligned and counted using Cell Ranger software (10x Genomics), which yielded feature-barcode matrices for each sample with each row of the matrix corresponding to a region of chromatin predicted to be open (also called a “peak”) and fragment files containing all sequenced fragments associated with each nucleus. Target regions were defined as transcription start sites and promoter, enhancer, and DNase hypersensitivity regions based on ENCODE (*26, 45*) and Ensembl 95 (*46*) annotations of these regulatory features.

The Signac platform version 0.2.5 (https://github.com/timoast/signac) was used to analyze these data in R. Nuclei that did not pass quality control metrics assessing nucleosome banding pattern (*47*), transcription start site (TSS) enrichment, total number and proportion of reads mapping to peaks, and proportion of reads falling in “blacklisted” regions (*26, 48*) were excluded from downstream analysis. Outlying nuclei were defined as follows: less than 3,000 or greater than 20,000 fragments mapping to peak regions (representing shallowly sequenced nuclei and multiplets); less than 15% reads mapping to peaks; more than 5% of reads mapping to blacklisted regions; nucleosome signal greater than 10, where nucleosome signal is equal to the number of fragments between 147-294 bp divided by fragments smaller than 147 bp; TSS enrichment score less than 2. TSS enrichment score is a measure of signal to noise ratio within a dataset, calculated as the center of the normalized distribution of reads mapping to TSSs (*26, 49*). Within the Signac package, term frequency-inverse document frequency (TF-IDF) normalization and dimensional reduction were completed for each of the four sample datasets individually. Linear dimensional reduction was accomplished by application of singular value decomposition (SVD) to perform a latent semantic indexing (LSI) transformation, a process analogous to principal component analysis.

The snATACseq datasets were merged after quantification of the peaks found in each sample individually relative to their combined set of peaks. Non-linear dimensional reduction using the UMAP technique and shared nearest neighbor graph-based clustering allowed identification of distinct cell populations as in snRNAseq analysis. A gene-activity matrix was created in Signac by examining the intersection between all fragments and coding regions of the genome including promoter regions, defined as the genomic regions specified by Ensembl (*46*) gene coordinates plus 2000 bp upstream of each gene. Differential gene activity around known cell type marker genes between clusters was used to assign each cell population to a human brain cell type, similar to the use of differential marker gene expression to categorize snRNAseq clusters. Cell type assignments were validated by transfer anchor-based integration (*36*) of the merged snATACseq dataset with a merged snRNAseq dataset derived from samples from the same four donors.

### Bulk ATACseq of iPSC-derived cells

For bulk ATAC sequencing of iMicroglia, live cells were collected on differentiation day 28 by FACS as previously described. Nuclei were extracted from 50,000 live iMicroglia and processed for library construction according to previously published protocols (*47, 50*). iFbN were collected at day 63 using accutase (StemCell Technologies #07920). Cells were counted using an automated cell counter (BioRad #1450102) and aliquoted 50,000 cells per sample.

Samples were done in triplicate and cells were lysed and transposed as previously published (*47, 50*). Illumina-compatible libraries were submitted to Psomagen or NISC for sequencing. The libraries were pooled and sequenced on an Illumina NovaSeq 6000 system to achieve a targeted sequencing depth of at least 25 million read pairs per sample. Adapter sequences were trimmed from the resulting DNA fragment sequences using NGmerge (*51*) and aligned to the GRCh38 (hg38) genome build with bowtie2 (*52*). Peaks were called using Genrich (https://github.com/jsh58/Genrich) after removal of PCR duplicates and mitochondrial reads. Peaks falling in blacklisted regions (*48*) were excluded.

### Integrated analysis of snATACseq and bulk ATACseq data

Integrated analysis of bulk ATACseq of iMicroglia and snATACseq of human frontal cortex was performed on the NIH HPC Biowulf cluster. To facilitate integration, nuclei barcodes with corresponding cell type assignments and sample ID were first extracted from the single nuclei dataset. Sinto (https://timoast.github.io/sinto/index.html) was then used to filter aligned fragments tagged by the barcodes of interest, producing separate BAM files for each cell type from each of the 4 donor datasets. These BAM files were merged to yield one BAM file containing all fragments associated with each cell type in the merged snATACseq dataset, similar to the output of bulk ATACseq. Peaks were called using Genrich as above for each brain cell type dataset. The cell type BAM files derived from snATACseq along with the BAM files resulting from bulk ATACseq of iPSC-derived cells were filtered to exclude mitochondrial reads, PCR duplicates, and non-unique alignments. The filtered BAM files were indexed, then DiffBind (*53*) was used to generate peak counts for each dataset. Finally, differential accessibility between datasets was assessed by DESeq2 (*54*) analysis.

### Genome editing of human iPS cells

CRISPR/Cas editing protocols were adapted from Skarnes *et al.* 2019 (*55*). Briefly, a ribonucleoprotein (RNP) consisting of complexed recombinant Cas enzyme and a chemically synthesized guide RNA (gRNA) were introduced into iPSCs by nucleofection along with a donor template single-stranded oligodeoxynucleotide (ssODN) containing the targeted single nucleotide substitution. The SNP rs76904798[C/T] was targeted with Cas12a on account of its proximity to the Cas12a protospacer adjacent motif *TTTN*. gRNA/donor template pairs were designed manually and confirmed using the CRISPOR (*56*) tool (http://crispor.tefor.net).

Recombinant Alt-R A.s. Cas12a (Cpf1) Nuclease Ultra (IDT #10001272), custom CRISPR-Cas12a crRNA (5’-ATTGTTACACGCTTTCCCAAA-3’), and custom donor template ssODN (5’-TGATAGTACCTTTTATATATTCTTCTTATTAATTTTTAATTGTTA**T**ACGCTTTCCCAAAATCTTCCAGCCTATGGTCATCTTTTAACTTTG-3’) were obtained from IDT.

iPSCs were cultured in E8 medium on matrigel-coated 6-well plates prior to editing experiments. Functional gRNA was prepared by reconstitution of the crRNA to a concentration of 100μM in Nuclease-free Duplex Buffer (IDT #11-01-03-01). The donor template ssODN was resuspended to a concentration of 200μM in DPBS. RNP complexes were formed by combining gRNA and Cas enzyme in PBS and incubating at room temperature for 30 minutes – 1 hour.

Meanwhile, iPSCs were rinsed with DPBS and incubated with TrypLE-Select (ThermoFisher #A1217701) for 2 minutes to achieve single cell dissociation. Cells were collected with DMEM/F12 and counted using a TC20 Automated Cell Counter (BioRad). Per editing reaction, 8 x 10^5^ cells were pelleted by centrifugation at 1000 RPM for 3 minutes. Donor template ssODN was added to the RNP complex immediately prior to nucleofection. Pelleted cells were resuspended in 100 μL of complete P3 Primary Cell Solution (Lonza #V4XP-3024), then mixed with the RNP/ssODN by trituration. The cell suspension was transferred to a Nucleocuvette (Lonza #V4XP-3024) and nucleofection was completed by electroporation using pulse code CA-137 on a 4D-Nucleofector X Unit (Lonza). Nucleofected cells were transferred to a new matrigel-coated well in a 6-well plate in 4mL of E8 medium containing 10μM ROCK inhibitor Y-27632 (StemCell Technologies #72307). Medium was replaced the following day with fresh E8, with 10μM ROCK inhibitor added if cell survival was poor.

E8 medium was replaced daily until edited iPSC pools reached 80-90% confluency. Cells were then split with TrypLE-Select, and 15-25,000 cells were seeded to a matrigel-coated 10 cm dish in E8/10μM ROCK inhibitor. Remaining cells were re-plated in matrigel-coated 6-well plates, and frozen in Synth-a-Freeze reagent (ThermoFisher #A1254201) when confluent.

Monoclonal cell colonies were grown from sparsely seeded single cells over 6-9 days of culture in E8 medium. Edited colonies were picked individually with a pipette tip under 4x magnification (Bioimager microscope) and re-plated in matrigel-coated 96-well plates in E8/10μM ROCK inhibitor. Colonies were cultured until most wells in the plate were confluent, typically 7-10 days, then split for 1) propagation and 2) screening by Sanger sequencing using a 3730*xl* DNA Analyzer (Applied Biosystems). Colonies positive for the target edit and without other changes in the immediate vicinity (within 300-400 bp) were expanded and re-sequenced. For each target edit, two colonies negative for the target edit were also expanded and re-sequenced to serve as a “non-edited” control (NEC) in later experiments.

Genomic DNA from clonal cell lines positive for the target edit as well as from the NEC cell lines was submitted to Psomagen, Inc. for targeted amplicon sequencing to confirm genotype. A 428 bp amplicon was produced using the following primer pair for PCR amplification: 5’-CCTCAAGTGATCCACCCACC-3’, 5’-TCAGGAAGCTTGAGGCAAGG-3’. Sequencing depth of at least 10,000 paired end reads was achieved on a MiSeq V3 system (300 PE, with multiplexing of amplicon libraries). After trimming of adapter sequences with NGmerge, sequences were mapped to the GRCh38 (hg38) genome build with bowtie2. Genotype at and around the position of interest was visualized and quantified using the Integrative Genomics Viewer (*57, 58*) developed by the Broad Institute (http://software.broadinstitute.org/software/igv/).

---

**Fig. S1.**
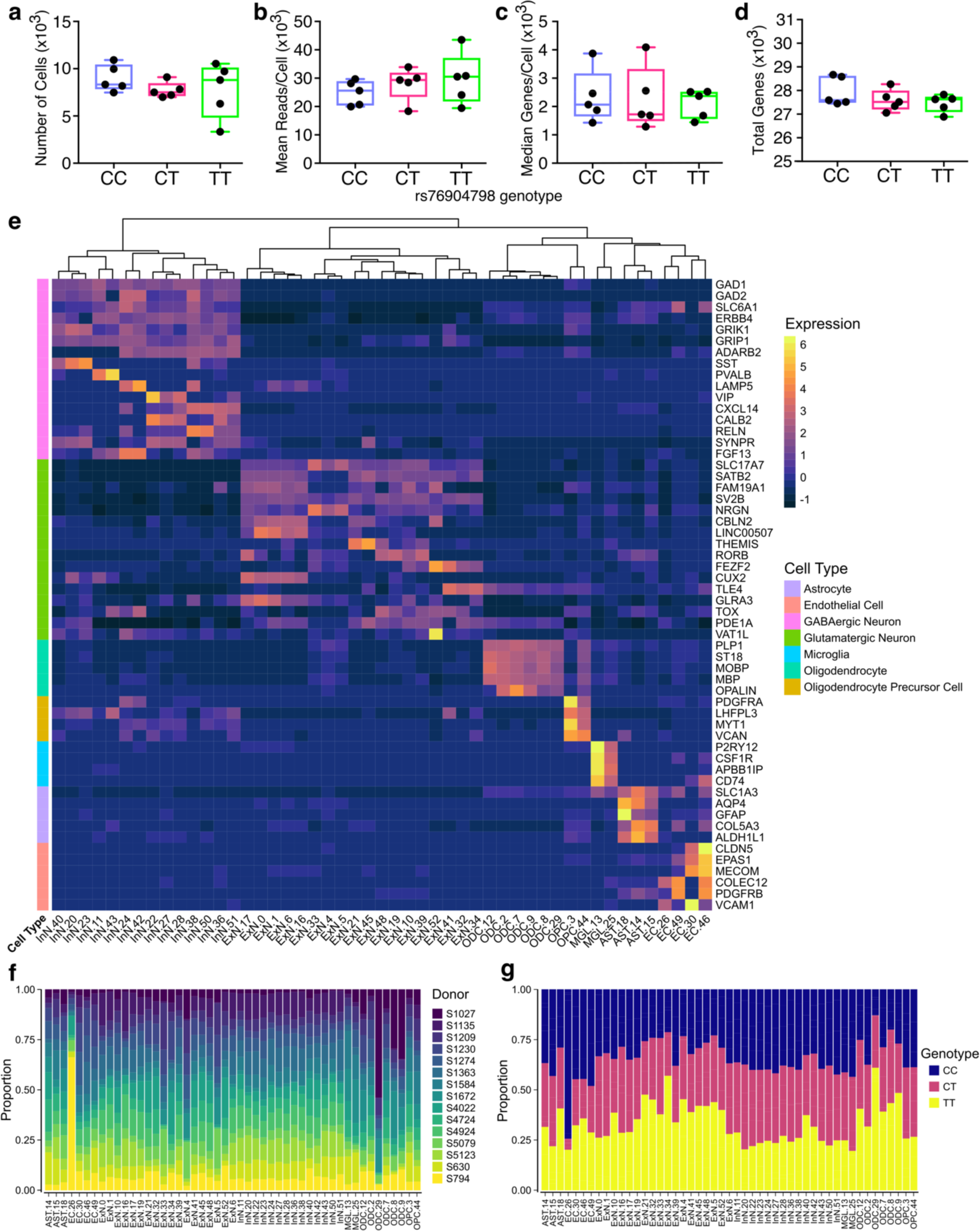
Box plots showing the distribution of (A) total number of cells measured, (B) average sequencing reads per cell, (C) median number of genes detected per cell, and (D) total number of genes detected per sample. There were no significant differences between genotype groups by one-way ANOVA with Tukey’s multiple comparisons test. (E) Heatmap of scaled expression of selected cell type marker genes, with a dendrogram indicating the relationship between cell populations using Ward’s hierarchical clustering method. Stacked bar charts showing the makeup of each cell population in terms of (F) donor and (G) rs76904798 genotype contribution.

**Fig. S2.**
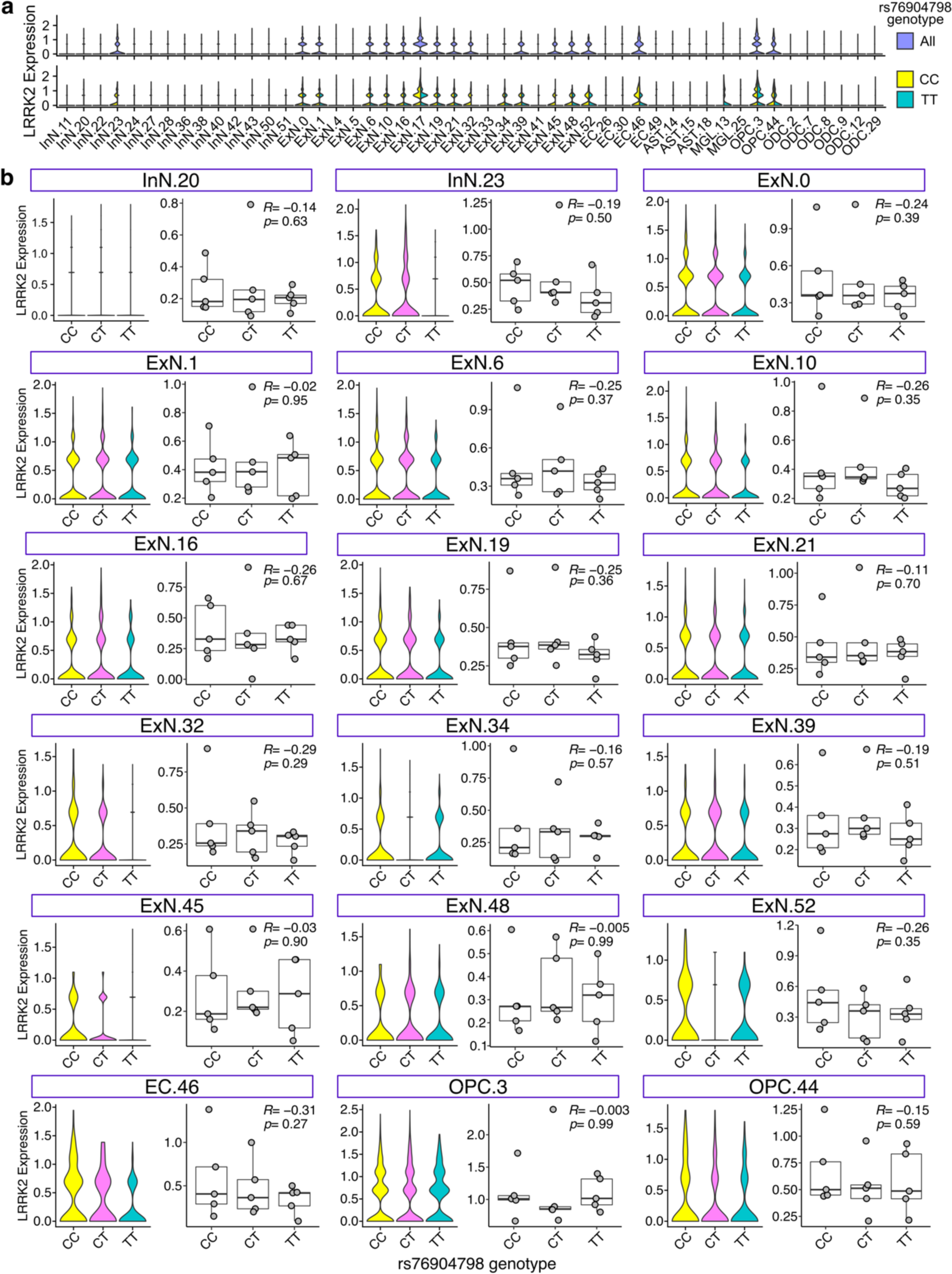
(A) Violin plots showing the distribution of *LRRK2* expression in each cell population overall, and split by genotype at rs76904798 (CC and TT only). (B) Violin plots split by rs76904798 genotype indicating *LRRK2* expression in each cell population in which *LRRK2* was detected in >18% of cells, as well as boxplots in which each point represents average *LRRK2* expression in cells contributed by each donor. The results of simple linear regression analysis of the correlation between number of minor alleles and *LRRK2* expression are shown (Pearson’s *R* and *p*-value, n = 15 within each cell population).

**Fig. S3.**
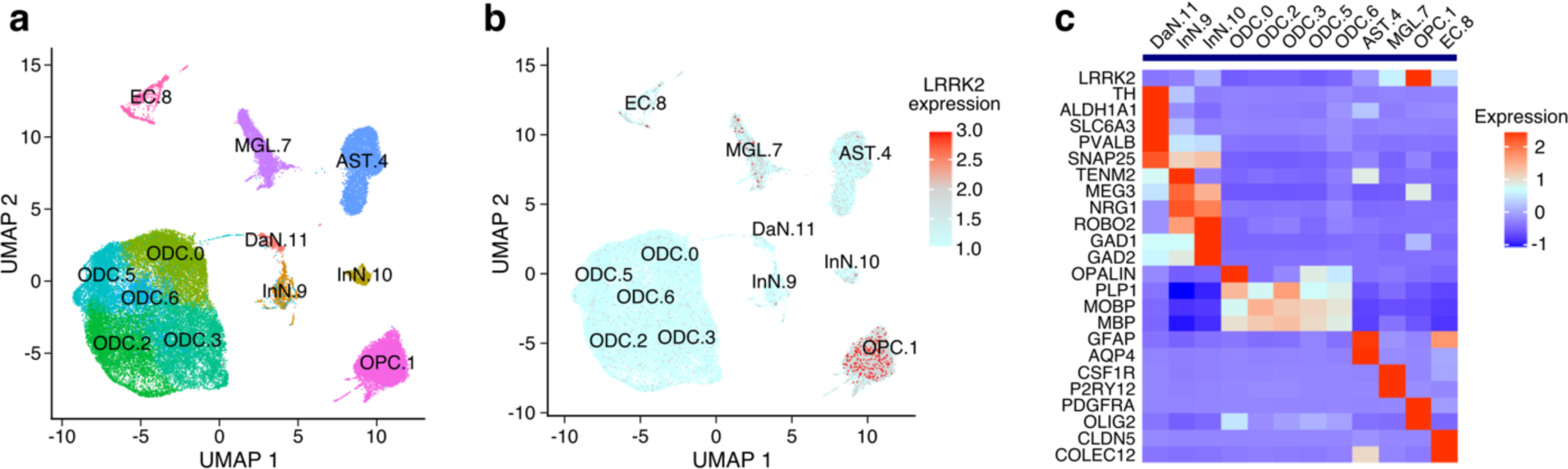
(A) UMAP plot of cell populations identified in human substantia nigra following integration and clustering analysis of snRNAseq datasets made publicly available by Welch *et al.* 2019^22^ and Agarwal *et al.* 2020^6^. A total of 46,109 nuclei are represented. (B) Distribution of *LRRK2* expression in the identified nigral cell populations. (C) Heatmap showing the average expression of known cell type marker genes along with *LRRK2* in each population.

**Fig. S4.**
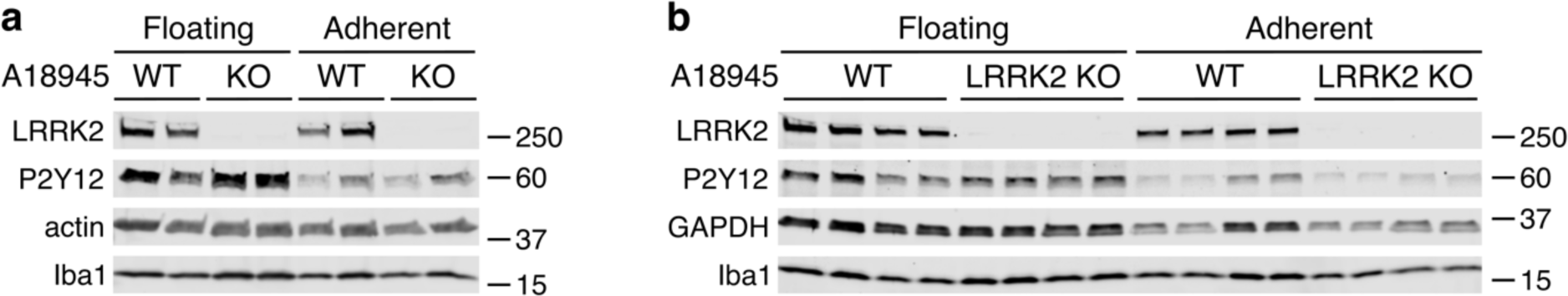
Western blot analysis of iMicroglia differentiated from A18945 WT and LRRK2 KO iPSC lines demonstrating LRRK2 expression in both floating and adherent cell populations, (A) technical n = 2 and (B) technical n = 4.

**Fig. S5.**
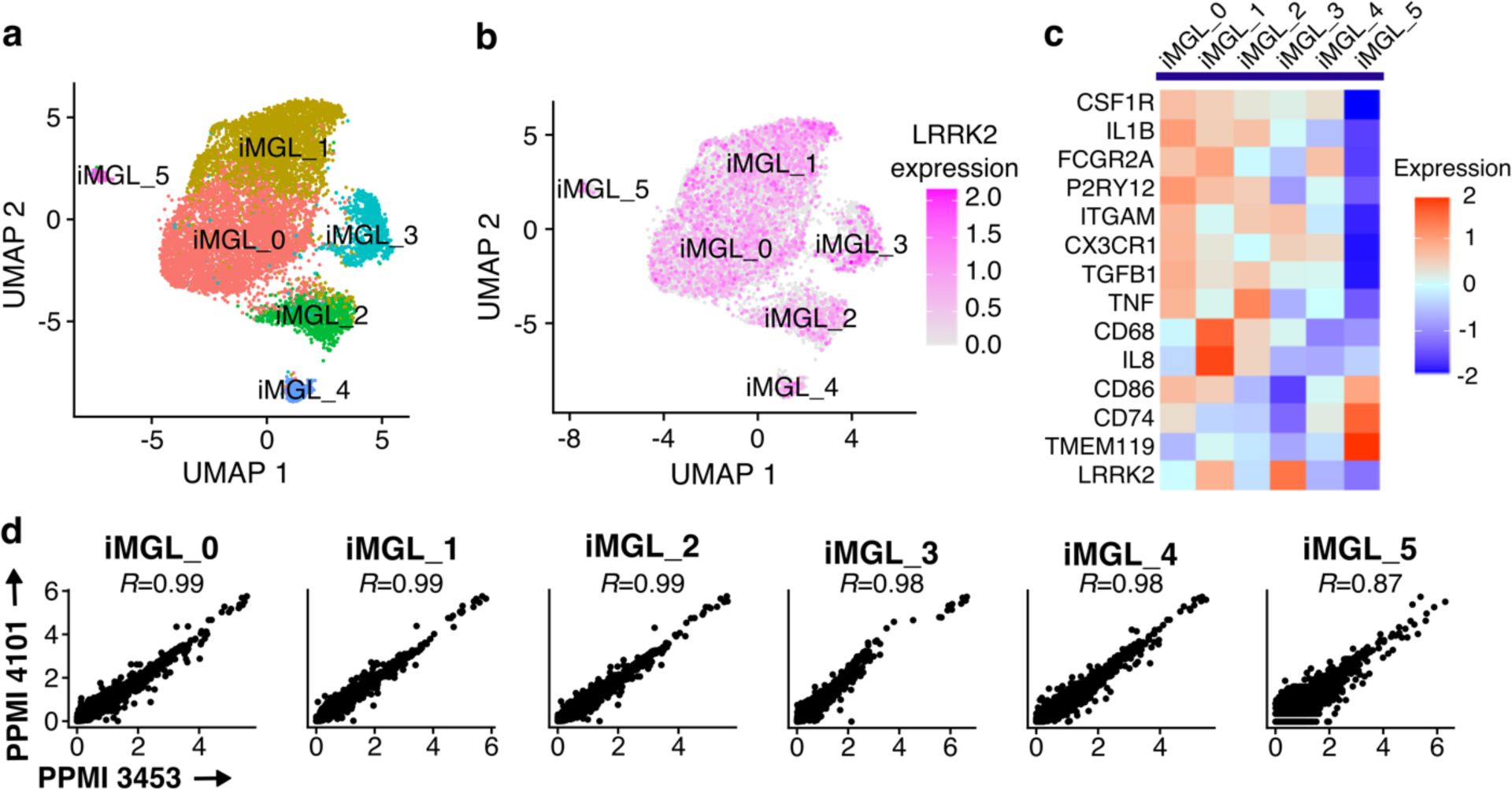
(A) UMAP visualization showing six cell populations identified by clustering analysis of scRNAseq of iMicroglia differentiated from two PPMI cell lines (3453 and 4101). (B) UMAP plot depicting the distribution of *LRRK2* expression. (C) Heatmap illustrating the relative expression of known microglia markers in each cell population. (D) Scatter plots describing the correlation between average gene expression in cells from each cell line in each cluster, described by Pearson’s *R*. For all correlation values, df = 17106, *p* < 2.22 x 10^-16^.

**Fig. S6.**
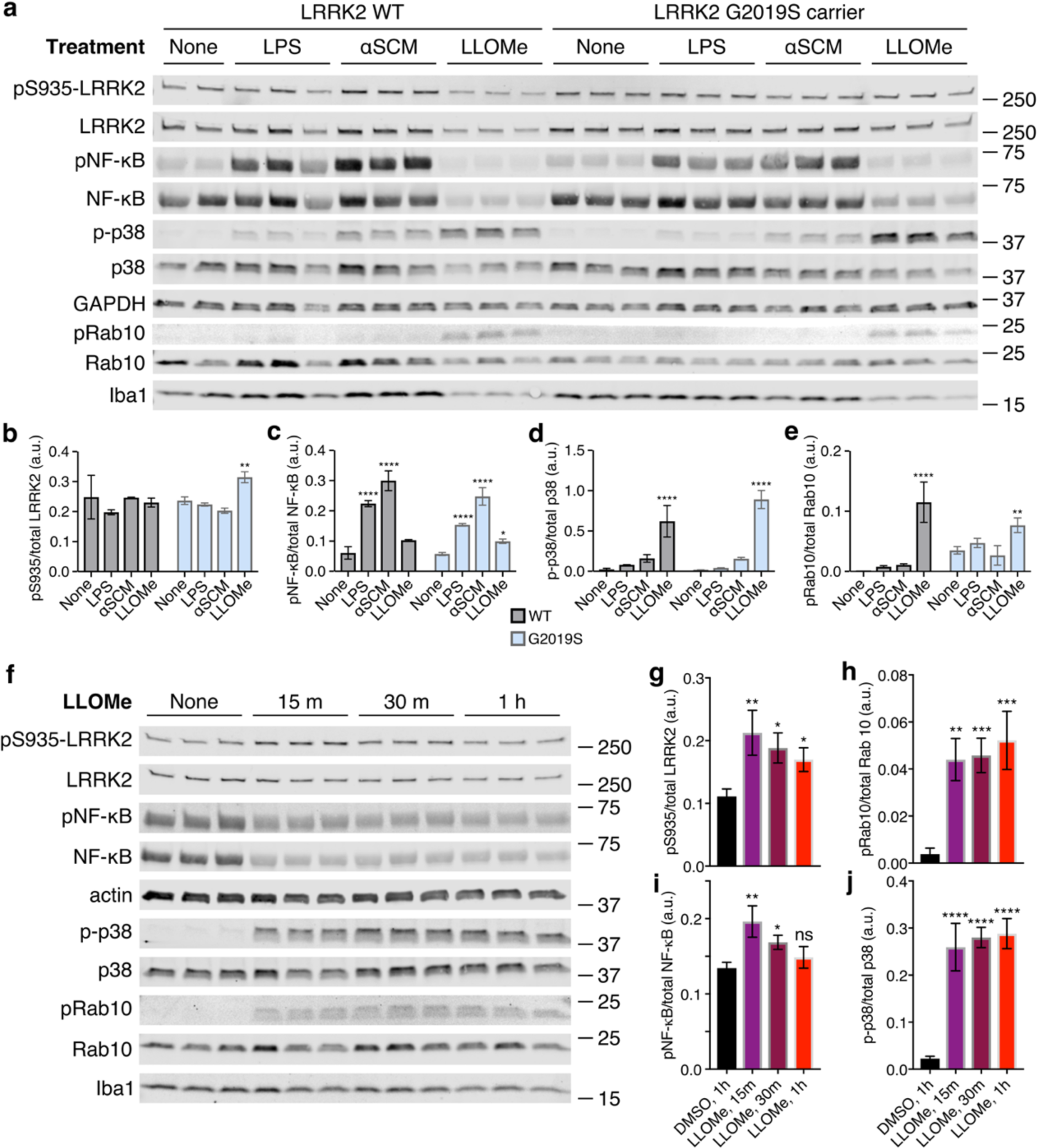
(A) Comparison of treatment conditions including 2-hour incubation with LPS (100ng/mL), α-synuclein conditioned medium (αSCM, 1x), or lysosomal inhibitor LLOMe (1mM) by western blot analysis of iMicroglia differentiated from two PPMI cell lines, one containing the LRRK2 kinase-activating mutation LRRK2-G2019S with *n* = 2-3 technical replicates. (B-E) Quantification of phosphorylated protein relative to the total protein amount. Statistical significance by two-way ANOVA with Dunnett’s multiple comparisons test is shown relative to the “None” un-treated control group for each cell line. (F) Time course study for optimization of LLOMe treatment time with iMicroglia differentiated from one PPMI cell line treated for 15 minutes, 30 minutes or 1 hour, with *n* = 3 technical replicates per time point. An equivalent volume of DMSO was added to untreated cells for 1 hour. (G-J) Quantification of phosphorylated protein relative to the total protein amount. The results of Dunnett’s multiple comparisons test following one-way ANOVA comparing LLOMe-treated cells to DMSO-treated control cells are shown; *P < 0.05, **P < 0.01, ***P < 0.001, ****P < 0.0001.

**Fig. S7.**
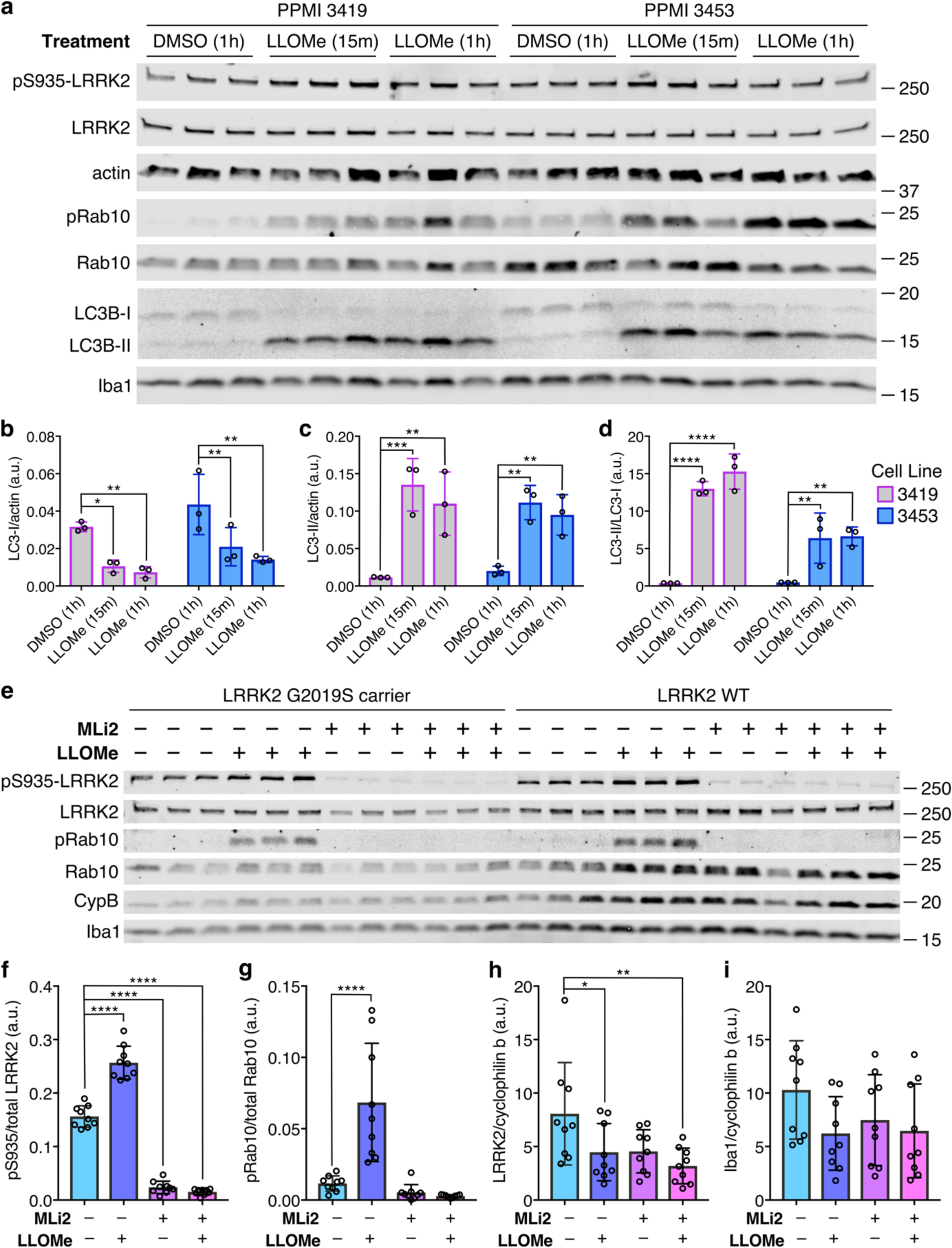
(A) Further characterization of iMicroglia response to LLOMe treatment times of either 15 minutes or 1 hour by immunoblot, using cells differentiated from two PPMI iPSC lines with technical *n* = 3. Untreated cells were incubated with an equivalent volume of DMSO for 1 hour. (B-D) Quantifications of LC3 levels, where decreased LC3-I and increased LC3-II are expected to correspond with upregulation of autophagy. Two-way ANOVA with Sidak’s multiple comparisons test were performed, and the results of comparisons between treated cells to untreated cells of the same cell line are shown; *P < 0.05, **P < 0.01, ***P < 0.001, ****P < 0.0001. (E) Representative western blot showing measurement of LRRK2 kinase dependency of LRRK2 kinase response to 15-minute LLOMe treatment, using iMicroglia differentiated from two PPMI iPSC lines (one heterozygous for LRRK2 kinase-activating mutation LRRK2-G2019S) with *n* = 3 technical replicates. LLOMe treatment assay was performed after 2-hour pretreatment of cells with LRRK2 kinase inhibitor MLi-2, or an equivalent volume of DMSO. The experiment was repeated with iMicroglia derived from a third PPMI iPSC line (WT-LRRK2), and the measurements from all three cell lines were aggregated for analysis. (F-I) Quantifications of assayed proteins (*N* = 3 cell lines, *n* = 9 with technical replicates). Statistically significant results of one-way ANOVA followed by Dunnett’s multiple comparisons test are shown; *P < 0.05, **P < 0.01, ***P < 0.001, ****P < 0.0001.

**Fig. S8.**
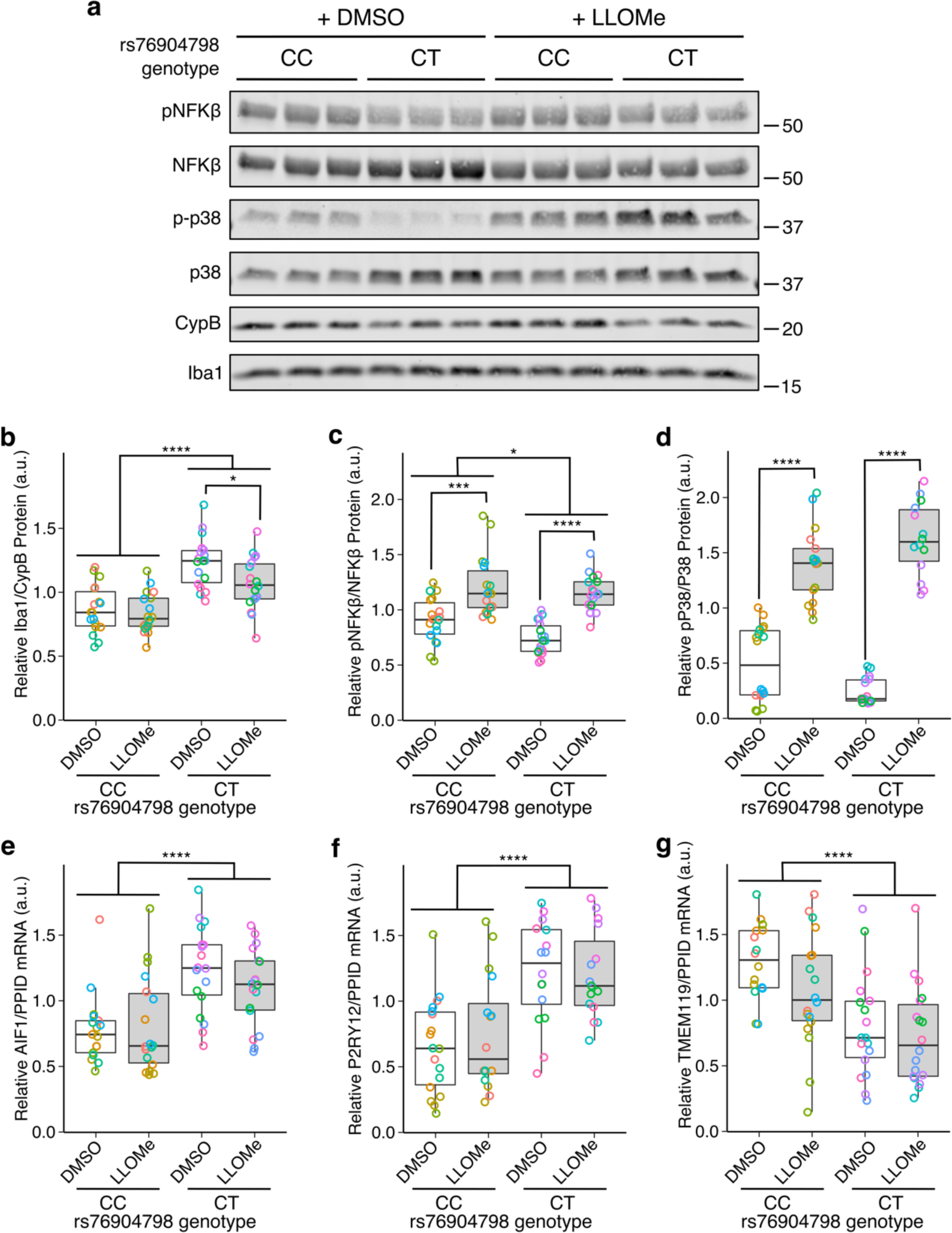
(A) Immunoblot of additional proteins in a representative western blot analysis of differentiated iMicroglia, PPMI 3411 (rs76904798-CC) and PPMI 3953 (rs76904798-CT), with immunoblotting for phospho-NF-κB, total NF-κB, phospho-p38, total p38, cyclophilin b (CypB) and Iba1 -/+ 15-minute LLOMe treatment. CypB and Iba1 bands are the same as those shown in Fig. 2c. Quantifications of normalized protein (B-D) and normalized mRNA (E-G) levels are presented relative to the mean value for each paired experiment. The *p*(genotype) results of two-way ANOVA are indicated where significant; *N* = 6 cell lines per genotype, *n* = 17-18 per genotype-treatment group with technical replicates. Significant results of Tukey’s multiple comparisons test between treatment groups are indicated.

**Fig. S9.**
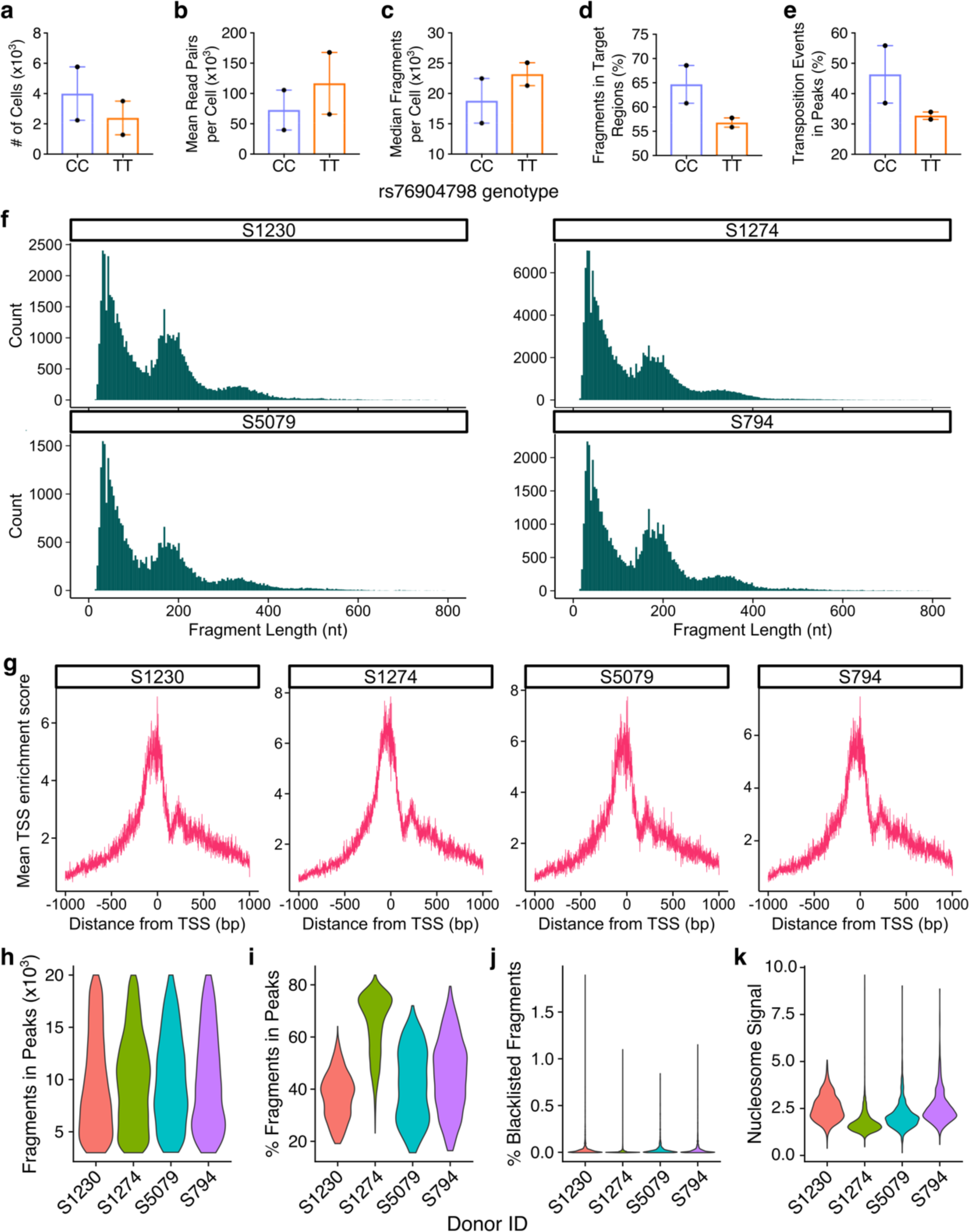
Bar graphs describing (A) total number of cells measured per sample (*p* = 0.52), (B) average sequencing depth per cell (*p* = 0.54), (C) median number of fragments mapped per cell (*p* = 0.40), (D) fraction of fragments mapping to regions of interest per sample (*p* = 0.19), and (E) proportion of Tn5 cut sites falling within called peaks per sample (*p* = 0.29). There were no significant differences in these parameters between rs76904798 genotype groups (unpaired Student’s *t* test, df = 2). Error bars represent SEM. (F) Histograms depicting the number of mapped reads of each length in nucleotides (nt) within each sample, labeled by donor ID. (G) Average transcription start site (TSS) enrichment score of nuclei within each sample, relative to distance in base pairs (bp) from annotated TSSs. (H-K) Violin plots showing the distribution of total number of fragments, percentage of fragments encompassing peaks, percentage of fragments encompassing blacklisted regions, and nucleosome signal, respectively, of nuclei contributed by each of the four samples in the merged dataset.

**Fig. S10.**
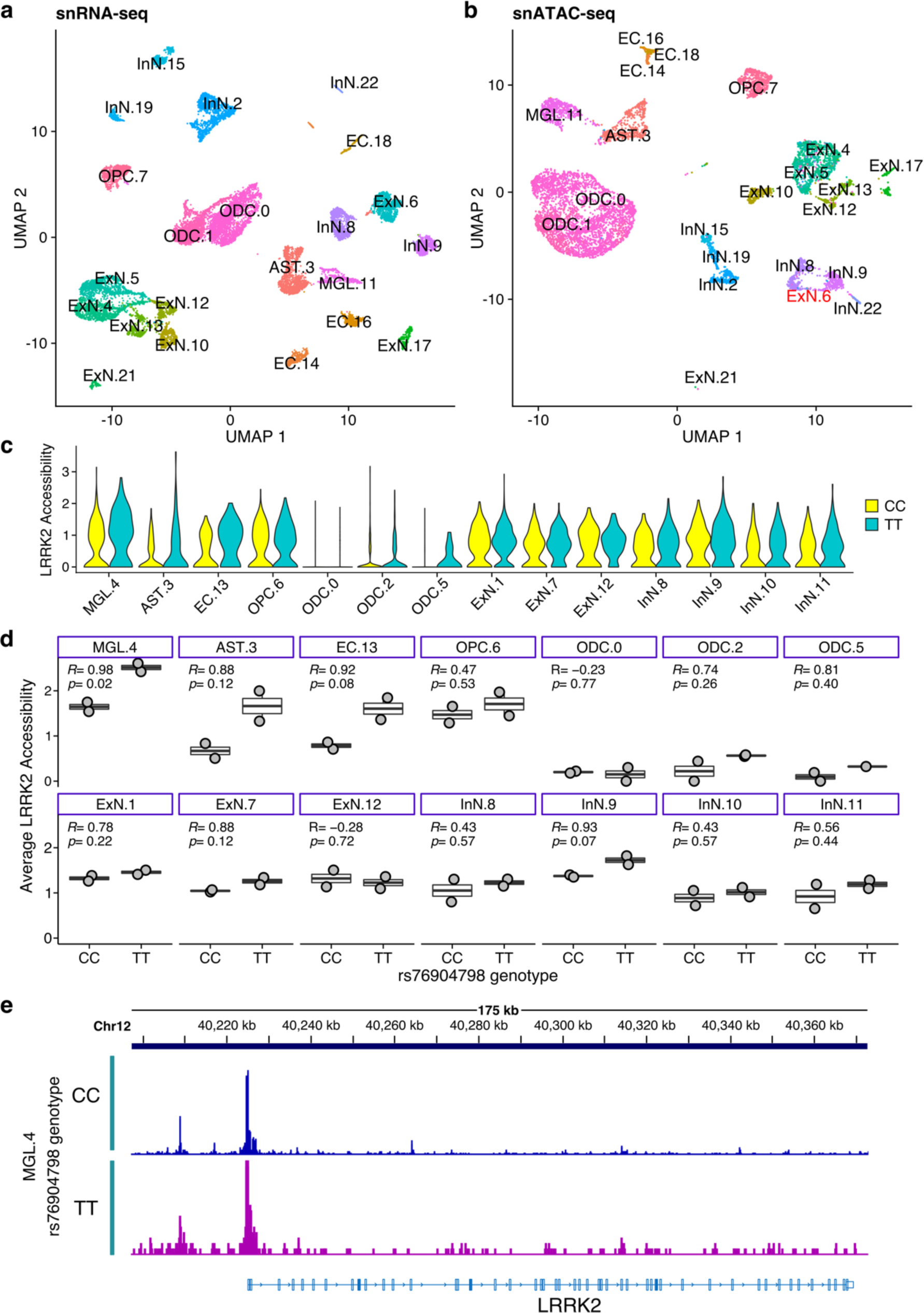
(A) UMAP visualization of cell populations identified by snRNAseq of 16,148 frontal cortex nuclei derived from the same four donors contributing to the snATACseq experiment. (B) UMAP plot of clusters distinguished by snATACseq analysis of 7,732 frontal cortex nuclei, labeled with the cell type assignment predicted by comparison with the snRNAseq analysis. (C) Split violin plot illustrating *LRRK2* gene accessibility in rs76904798-CC cells vs. rs76904798-TT cells of each population. (D) Boxplots showing average *LRRK2* gene accessibility in cells contributed by each donor. Pearson’s *R* and *p*-values describing the correlation between number of minor alleles and *LRRK2* gene accessibility derived by simple linear regression analysis are shown (*n* = 4 within each cell population). (E) IGV browser view of *LRRK2* gene accessibility in CC vs TT cells of frontal cortex snATACseq population “MGL.4”.

**Fig. S11.**
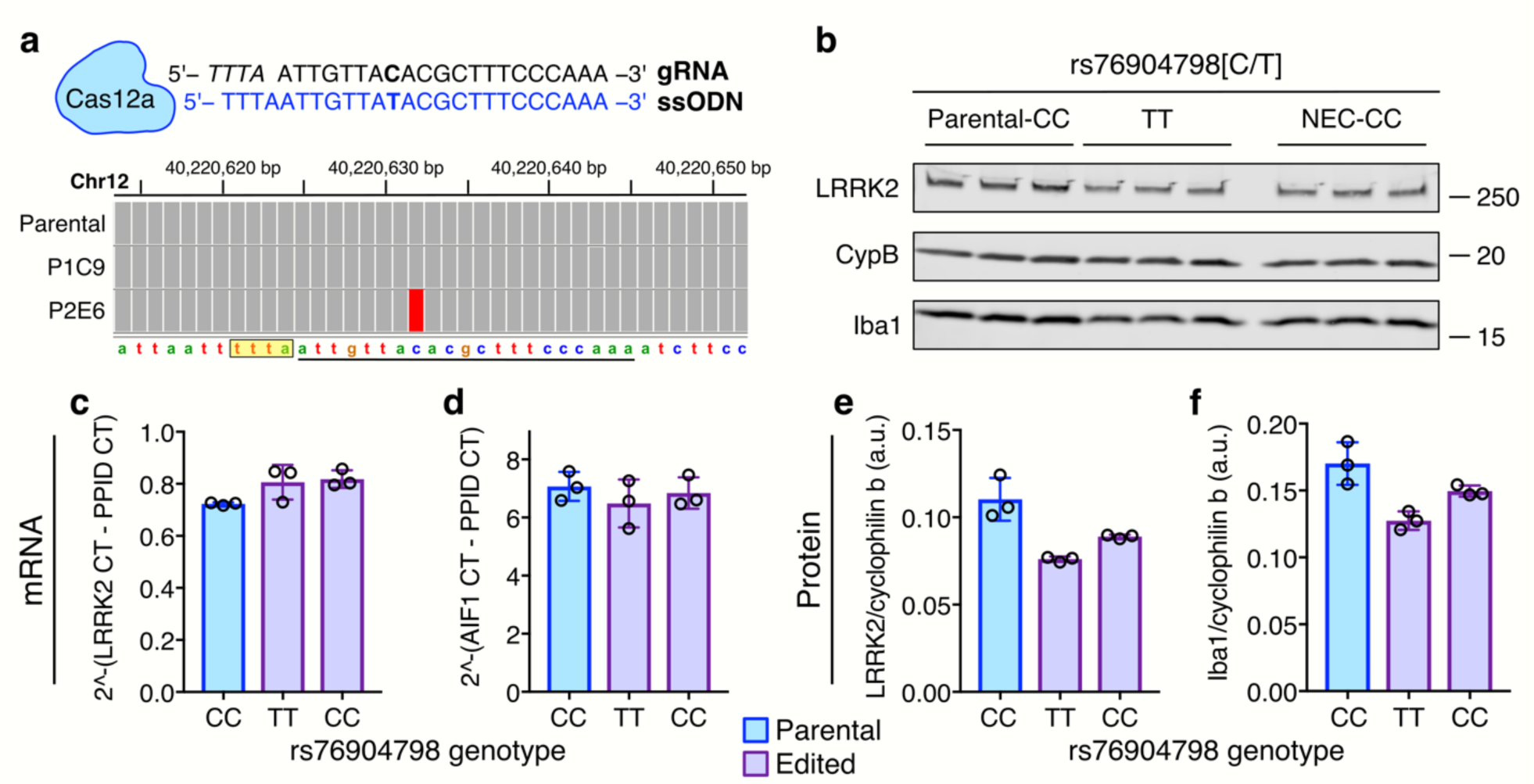
The rs76904798 variant itself does not underlie PD risk association. (A) Depiction of the gRNA and donor template used to make a Cas12a-directed single nucleotide substitution in PPMI 3471 iPSC line (genotype rs76904798-CC), to produce an edited iPSC line with genotype rs76904798-TT as confirmed by targeted amplicon sequencing. An IGV browser view of the sequencing results are shown for the original PPMI cell line (“Parental”), a non-edited control (NEC) clone (“P1C9”), and the correctly edited clone (“P2E6”) used in differentiation experiments. (B) Western blot analysis of iMicroglia differentiated from the parental cell line (rs76904798-CC), the edited cell line (rs76904798-TT), and the NEC line (rs76904798-CC). Normalized quantifications of *LRRK2* (C) and *AIF1* (D) mRNA and LRRK2 (E) and Iba1 (F) protein levels, technical *n* = 3. No significant differences in these measurements were detected by one-way ANOVA followed by Dunnett’s multiple comparisons test.

**Fig. S12.**
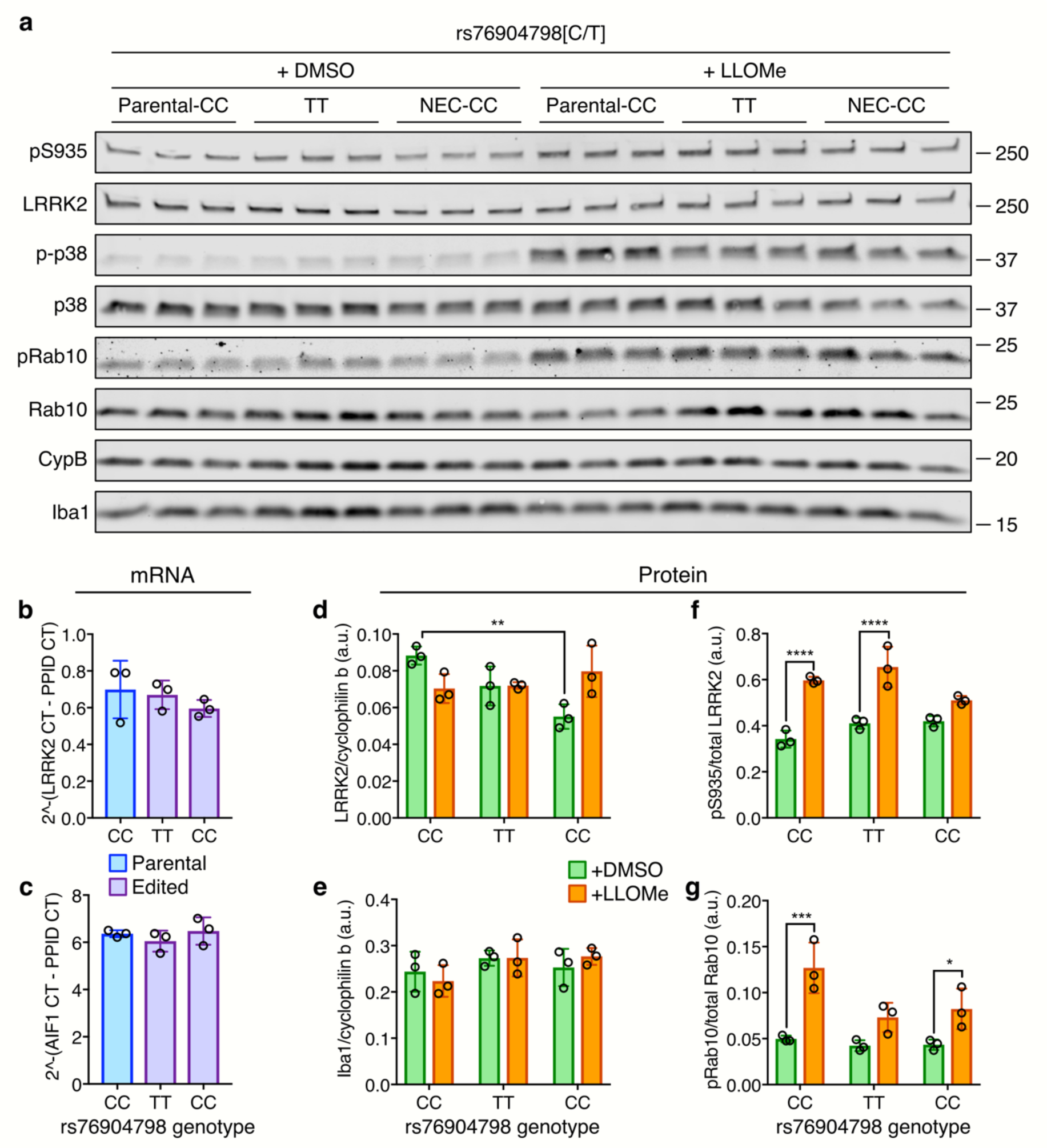
(A) Western blot analysis of iMicroglia differentiated from the parental PPMI 3471 iPSC line (rs76904798-CC), edited clone 3471-2E6 (rs76904798-TT), and NEC clone 3471-1C9 (rs76904798-CC) following 15-minute treatment with 1mM LLOMe. Results of immunoblotting for phospho-Ser^935^-LRRK2, total LRRK2, phospho-p38, total p38, phospho-Rab10, total Rab10, cyclophilin b (CypB) and Iba1 are shown. (B-G) Quantifications of normalized mRNA or protein levels, technical n = 3. The results of two-way ANOVA followed by Sidak’s multiple comparisons test are indicated where statistically significant; *P<0.05, **P<0.01, ***P<0.001, ****P<0.0001.

**Table S1.**
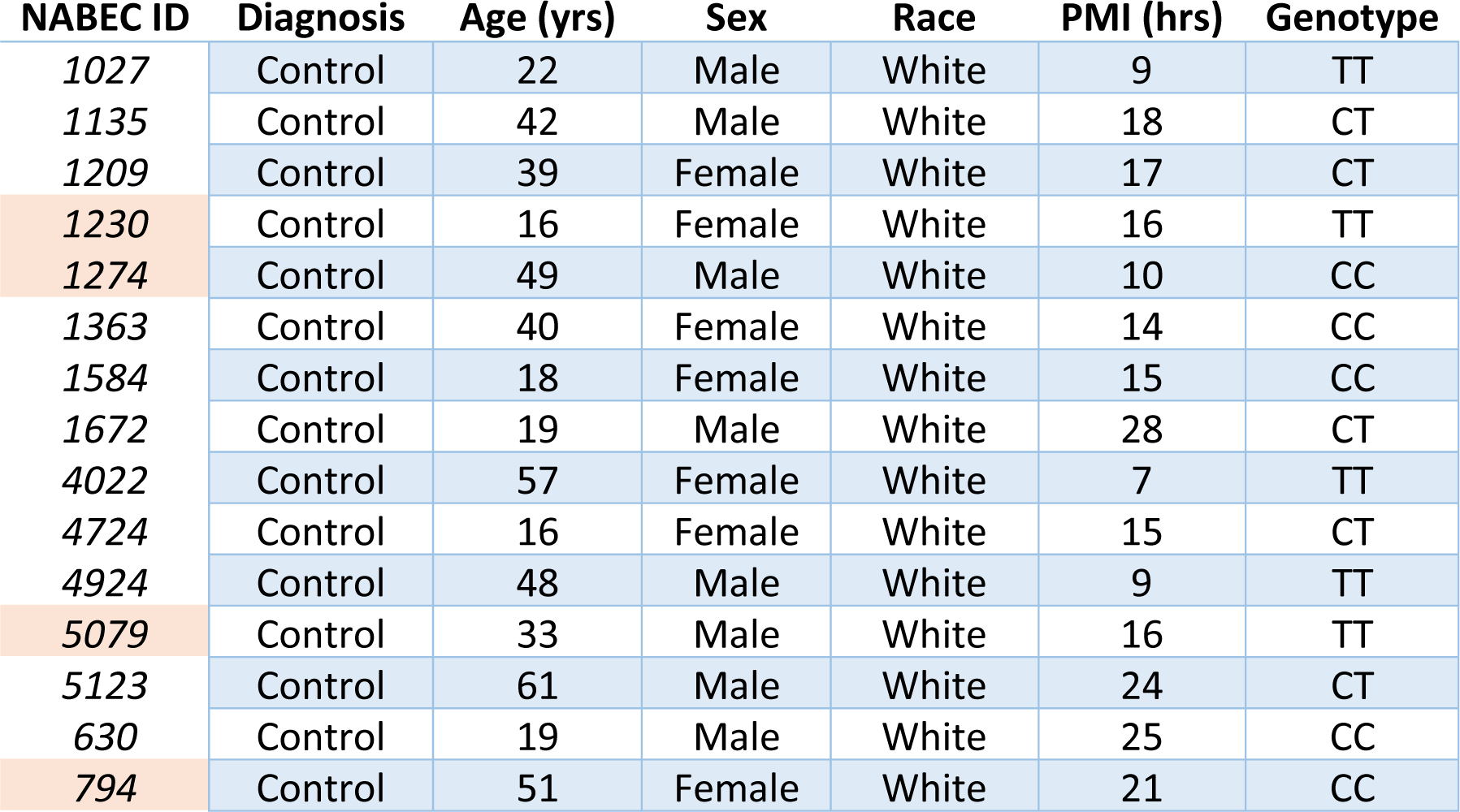
Demographic description of human frontal cortex tissue donors. NABEC = North American Brain Expression Consortium. PMI = Postmortem Interval. Genotype at rs76904798[C/T]. Red-shaded cells indicate donors from whom samples were submitted to both single nuclei RNA and single nuclei ATAC sequencing.

**Table S2.** Characteristics of PBMC donors from which iPSC lines were derived. Shading indicates pairing of cell lines for differentiation. PPMI = Parkinson’s Progression Markers Initiative. PD = Parkinson’s Disease; HC = Healthy Control. Genotype at rs76904798[C/T].

| PPMI ID | Diagnosis | Birth Year | Sex | Ethnicity | Genotype |
| --- | --- | --- | --- | --- | --- |
| 3446 | PD | 1936 | Male | Caucasian | CT |
| 3471 | PD | 1938 | Male | Caucasian | CC |
| 3467 | PD | 1945 | Male | Caucasian | CT |
| 3234 | PD | 1943 | Male | Caucasian | CC |
| 3473 | PD | 1957 | Female | Caucasian | CT |
| 3186 | PD | 1951 | Female | Caucasian | CC |
| 3664 | PD | 1951 | Female | Caucasian | CT |
| 3453 | HC | 1951 | Female | Caucasian | CC |
| 3953 | PD | 1958 | Male | Caucasian | CT |
| 3411 | HC | 1970 | Male | Caucasian | CC |
| 4101 | PD | 1946 | Female | Caucasian | CT |
| 3419 | PD | 1943 | Female | Caucasian | CC |

## Notes

### Competing Interest Statement

The authors have declared no competing interest.

